# Deep evolutionary origin of gamete-directed zygote activation by KNOX/BELL transcription factors in green plants

**DOI:** 10.1101/2020.04.08.031930

**Authors:** Tetsuya Hisanaga, Shota Fujimoto, Yihui Cui, Katsutoshi Sato, Ryosuke Sano, Shohei Yamaoka, Takayuki Kohchi, Frédéric Berger, Keiji Nakajima

## Abstract

KNOX and BELL transcription factors regulate distinct steps of diploid development in the green lineages. In the green alga *Chlamydomonas reinhardtii,* KNOX and BELL proteins are inherited by gametes of the opposite mating types, and heterodimerize in zygotes to activate diploid development. By contrast, in land plants such as Physcomitrella and Arabidopsis, KNOX and BELL proteins function in meristem maintenance and organogenesis during the later stages of diploid development. However, whether the contrasting functions of KNOX and BELL were acquired independently in algae and land plants is currently unknown. Here we show that in the basal land plant species *Marchantia polymorpha*, gamete-expressed *KNOX* and *BELL* are required to initiate zygotic development by promoting nuclear fusion in a manner strikingly similar to that of *C. reinhardtii*. Our results indicate that zygote activation is the ancestral role of KNOX/BELL transcription factors, which shifted toward meristem maintenance as land plants evolved.

## Introduction

The life cycles of eukaryotes alternate between diploid (2n) and haploid (n) phases through meiosis and fertilization (Bowman et al., 2016). In land plants, both the haploid and diploid phases are multicellular, producing gametophytic and sporophytic bodies, respectively. In bryophytes including liverworts, mosses, and hornworts, gametophytes are larger and more morphologically complex than sporophytes, which consist of only a few cell types. During the course of land plant evolution, the life cycle shifted toward a sporophyte-dominant style, presumably to facilitate adaptation to terrestrial environments where it is advantageous to generate large sporophytes that produce many spores. Consequently, the sporophytes of extant flowering plants (angiosperms) exhibit far more complex morphologies than their male and female gametophytes, the pollen grain and embryo sac, respectively, which are composed of only a few cells. This evolutionary transition in life cycle is thought to have been facilitated by the cooption of genes and/or gene regulatory networks that regulate gametophyte development to function in sporophyte development (Bowman, 2019). However, key steps of life cycle progression per se have continued to be driven by conserved regulators during land plant evolution, as recently reported for gametophytic sexual differentiation and gamete formation (Koi et al., 2016, Rövekamp et al., 2016, Higo et al., 2018, Yamaoka et al., 2018, Hisanaga et al., 2019a, Hisanaga et al., 2019b).

Homeodomain transcription factors (HD TFs) are developmental regulators that are evolutionarily conserved in eukaryotes. HD TFs are classified into two families, Three-Amino-Acid-Loop-Extension (TALE) and non-TALE, based on amino acid sequence similarity in the HD domain (Bertolino et al., 1995, Derelle et al., 2007). In the fungi *Saccharomyces cerevisiae* and *Coprinopsis cinerea*, TALE and non-TALE HD TFs are expressed in haploid cells of opposite mating types. These proteins heterodimerize in zygotes to regulate the expression of genes promoting the haloid-to-diploid transition (Goutte and Johnson, 1988, Herskowitz, 1989, Kües et al., 1992, Spit et al., 1998). In green plants, TALE HD TFs have diversified into the KNOX (KNOTTED1-LIKE HOMEOBOX) and BELL (BELL-LIKE) subfamilies. In the unicellular green alga *C. reinhardtii*, the KNOX protein GAMETE SPECIFIC MINUS1 (GSM1) and the BELL protein GAMETE SPECIFIC PLUS1 (GSP1) accumulate in the cytosol of *minus* and *plus* gametes, respectively. Upon fertilization, the two proteins heterodimerize and translocate to both male and female pronuclei to activate the expression of early zygote-specific genes. Loss-of-function mutations in either *GSP1* or *GSM1* result in pleiotropic phenotypes involving cellular rearrangements in zygotes, such as the loss of nuclear and mitochondrial fusion, lack of selective degradation of *minus*-derived chloroplast DNA and chloroplast membrane fusion, and defects in flagella resorption (Joo et al., 2017, Kariyawasam et al., 2019, Lee et al., 2008, Lopez et al., 2015, Nishimura et al., 2012).

In land plants, KNOX proteins are further diversified into class I (KNOX1) and class II (KNOX2) (Kerstetter et al., 1994, Mukherjee et al., 2009). The developmental functions of these proteins have been studied extensively in angiosperms such as maize (*Zea mays*), rice (*Oryza sativa*), and Arabidopsis (*Arabidopsis thaliana*) (reviewed in Hay and Tsiantis, 2010). Based on their expression patterns and the phenotypes of both knockout and overexpression lines, KNOX1 proteins are thought to promote cell proliferation in the meristematic tissues of aerial organs. The biological functions of *KNOX2* genes are somewhat elusive, but they are thought to act antagonistically to *KNOX1* to promote cell differentiation (Furumizu et al., 2015).

The apparent functional dissimilarity of KNOX proteins between *C. reinhardtii* and Arabidopsis (zygote activation versus cellular proliferation/differentiation) may reflect the large phylogenetic distance between these two species, as they separated into two major green plant lineages, Chlorophyta and Streptophyta, some 700 million years ago (Becker, 2013). Functional analyses of *KNOX* genes in the moss *Physcomitrella patens,* however, pointed to some commonality between the two *KNOX* functions in sporophyte development. The moss genome contains three *KNOX1* and two *KNOX2* genes, which are all primarily expressed in sporophytes, though expression of one *KNOX1* and two *KNOX2* genes is additionally detected in egg cells (Horst et al., 2016, Sakakibara et al., 2013, Sakakibara et al., 2008). A triple loss-of-function mutant of all three *KNOX1* genes was defective in cell division and differentiation in sporophytes, as well as spore formation (Sakakibara et al., 2008). By contrast, simultaneous knockout of the two *KNOX2* genes resulted in ectopic gametophyte formation in sporophyte bodies (Sakakibara et al., 2013). Thus, at least in one bryophyte species, KNOX1 and KNOX2 control sporophyte development via two pathways, with one ensuring proper sporophyte development (like *C. reinhardtii* GSM1) and the other promoting cell proliferation (like Arabidopsis KNOX1 proteins). While the transition of the role of KNOX/BELL from zygote activation to sporophyte morphogenesis likely arose during plant evolution, the point in plant phylogeny at which this transition occurred is unclear.

Here, we analyzed the roles of KNOX1 and BELL in the liverwort *Marchantia polymorpha*, a model species suitable to study evolution of sexual reproduction in plants (Hisanaga et al., 2019b). We uncovered unexpected conservation of KNOX/BELL function between the phylogenetically distant green plants *M. polymorpha* and *C. reinhardtii*, but not between the more closely related *M. polymorpha* and *P. patens*. Thus, the functional transition of KNOX/BELL from zygote activation to sporophyte morphogenesis occurred at least once in the land plant lineage independently of the acquisition of multicellular sporophytes. Additionally, we uncovered inverted sex-specific expression patterns of *KNOX* and *BELL* genes between *C. reinhardtii* and *M. polymorpha*, suggesting that anisogamy evolved independently of *KNOX/BELL* expression in gametes.

## Results

### Mp*KNOX1* is as an egg-specific gene in *M. polymorpha*

We previously reported that an RWP-RK TF MpRKD promotes egg cell differentiation in *M. polymorpha*. Loss-of-function Mp*rkd* mutant females grow normally and produce archegonia like the wild type, but their egg cells do not mature, instead degenerating after ectopic cell division and vacuolization (Koi et al., 2016). We made use of this egg-specific defect in Mp*rkd* to identify genes preferentially expressed in egg cells of *M. polymorpha*. Briefly, we collected ∼2,000 archegonia from two independent Mp*rkd* female mutant lines (Mp*rkd-1* and Mp*rkd-3*, Koi et al., 2016), each in two replicates. As a control, ∼4,000 archegonia were collected from wild-type females in four replicates. We extracted RNA from each pool and analyzed it by next-generation sequencing. Comparative transcriptome analysis identified 1,583 genes with significantly reduced mRNA levels in Mp*rkd* compared to wild-type archegonia (>3-fold and *false discovery rate* < 0.01; Figure 1A and Figure1 – figure supplement 1A). Among these, Mp*KNOX1* (Mp5g01600), the only class I *KNOX* gene in *M. polymorpha* (Bowman et al., 2017, Frangedakis et al., 2017), was strongly downregulated in Mp*rkd* vs. the wild type (∼500-fold) (Figure 1B). The MpKNOX1 polypeptide contains KNOX I, KNOX II, ELK, and Homeobox domains, as do KNOX proteins from green algae, mosses, ferns and flowering plants (Figure 1C).

**Figure 1.**
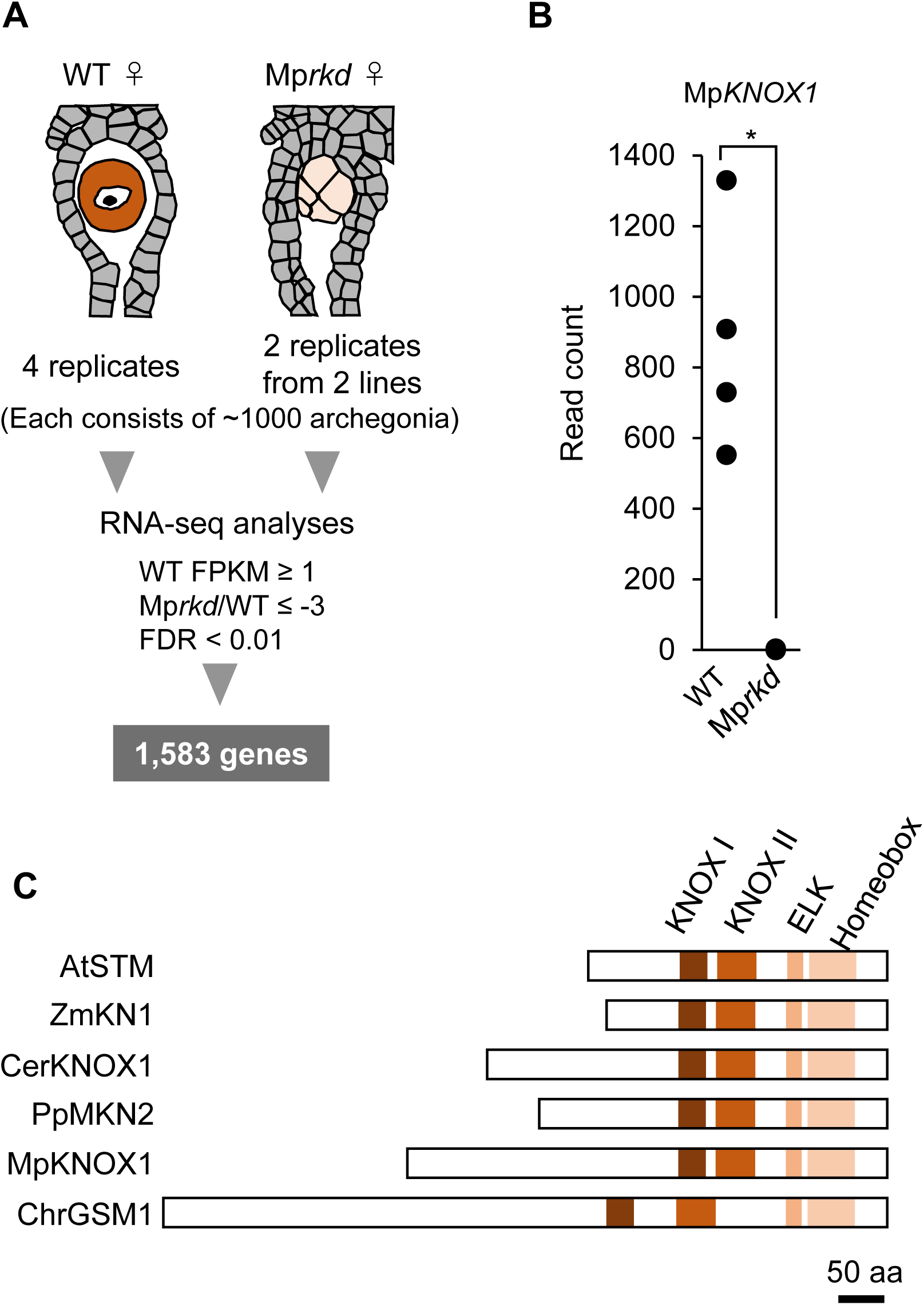
Comparative transcriptome analysis of wild-type and Mp*rkd* archegonia and identification of Mp*KNOX1* as an egg-specific gene. **A.** Schematic illustration of RNA-seq analysis comparing the archegonia transcriptomes from wild-type females and egg-deficient Mp*rkd* mutant females. About 4,000 and 2,000 archegonia were collected from wild-type and each of the two Mp*rkd* mutant lines, and randomly allocated into four and two replicates, respectively. **B.** A graph showing Mp*KNOX1* read counts in wild-type and Mp*rkd* archegonia. Asterisk indicates significant difference (*q* = 5.84E-83, *exactTest* in edgeR package). Not that four data points are overlapping in Mp*rkd*. **C.** Comparison of the domain arrangements of MpKNOX1 vs. representative class I KNOX proteins from *Arabidopsis thaliana* (AtSTM; GenBank accession number AEE33958.1), *Zea mays* (ZmKN1; AAP21616.1), *Ceratopteris richardii* (CerKNOX1, BAB18582.1), *Physcomitrella patens* (PpMKN2; AAK61308.2) and *Chlamydomonas reinhardtii* (ChrGSM1; ABJ15867.1).

Previous RNA-sequencing data (Bowman et al., 2017) indicate that Mp*KNOX1* is specifically expressed in female plants (Figure 1 – figure supplement 1B). To obtain the detailed expression patterns of Mp*KNOX1*, we performed RT-PCR analysis using RNA extracted from vegetative and reproductive organs of male and female gametophytes, as well as three-week-old sporophytes, and confirmed the specific expression of Mp*KNOX1* in archegoniophores (Figure 2A). No expression was detected in female or male thalli (leaf-like vegetative organs), antheridiophores (male reproductive branches) or sporophytes (Figure 2A). To visualize the cell- and tissue-specific expression patterns of Mp*KNOX1*, we generated a Mp*KNOX1* transcriptional reporter line (*MpKNOX1pro:H2B-GFP*). Consistent with the dramatically reduced Mp*KNOX1* transcript levels in egg-deficient Mp*rkd* archegonia (Figure 1B), reporter expression was specifically detected in egg cells (Figure 2C) and not in developing archegonia before the formation of egg progenitors (Figure 2B). Together, these data indicate that Mp*KNOX1* is an egg-specific gene in *M. polymorpha*.

**Figure 2.**
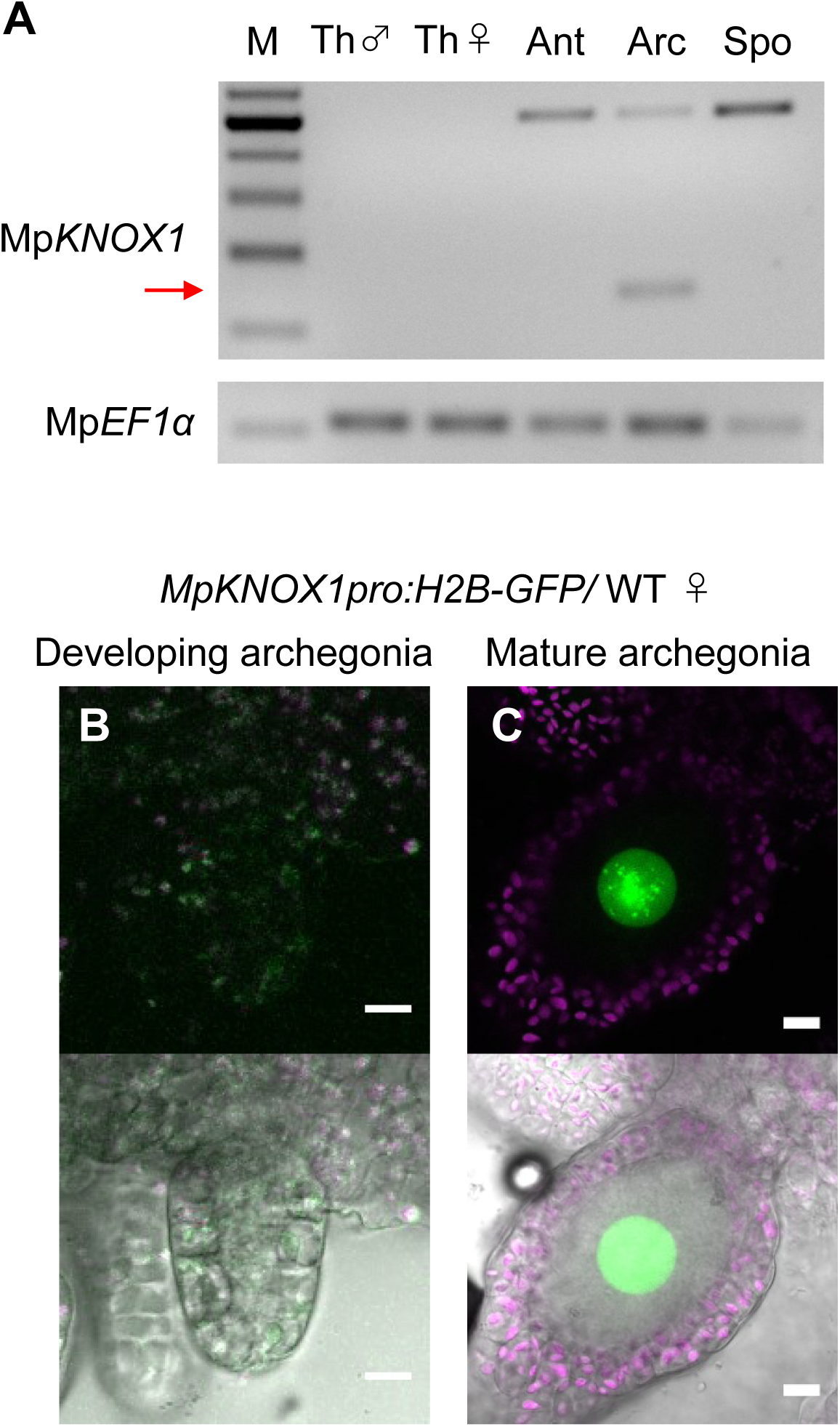
Mp*KNOX1* is specifically expressed in egg cells. **A.** RT-PCR analysis of Mp*KNOX1*. Lanes are labeled as follows: M, size markers, Th ♂, male thalli, Th ♀, female thalli, Ant, Antheridiophores, Arc, Archegoniophores, Spo, sporophytes of 3-week-old plants. Constitutively expressed Mp*EF1α* was used as a control. Red arrow indicates the expected size of PCR products from spliced Mp*KNOX1* mRNA. Bands at the top of the gel likely correspond to unspliced Mp*KNOX1* transcripts. Shown is a representative result from the experiments using three independently collected plant samples each with two technical replicates (two PCRs from each cDNA pool). **B and C.** Expression of the Mp*KNOX1* transcriptional reporter. Magenta, chlorophyll autofluorescence; green, GFP fluorescence. Lower panels are merged photographs of fluorescence and bright-field images. Bars, 10 μm.

### Zygote activation of wild-type *M. polymorpha* lags during karyogamy

The egg-specific expression of Mp*KNOX1* attracted our attention, as this pattern is in contrast to the previously reported function of KNOX1s in sporophyte morphogenesis in land plants (Furumizu et al., 2015, Sakakibara et al., 2008). Instead, the egg-specific expression of Mp*KNOX1* is reminiscent of *GSM1,* a KNOX homolog in *C. reinhardtii* that is expressed in *plus* gametes and activates zygote development after fertilization (Joo et al., 2017, Kariyawasam et al., 2019, Lee et al., 2008, Lopez et al., 2015, Ning et al., 2013, Nishimura et al., 2012). Therefore, we explored whether Mp*KNOX1* plays a role in zygote activation in *M. polymorpha*.

While the processes of gametogenesis and embryo patterning in *M. polymorpha* (Figure 3A) have been characterized histologically (Durand, 1908, Higo et al., 2016, Koi et al., 2016, Shimamura, 2016, Zinsmeister and Carothers, 1974), the subcellular dynamics associated with zygote activation have not been described in detail. To visualize these dynamics, we established a simple *in vitro* fertilization method for *M. polymorpha*. Briefly, several antheridiophores and archegoniophores were co-cultured in a plastic tube containing an aliquot of water for 1 h to allow sperm to be released from the antheridia and enter into the archegonia (Figure 3B). At the end of the co-culture period, sperm nuclei were visible in most eggs (Figure 3C), indicating that fertilization had been completed during the 1 h of co-culturing. Subsequently, sperm-containing archegonia were transferred to a fresh tube containing water and cultured for two weeks (Figure 3B). This experimental regime restricted the timing of sperm entry to the 1-h time window of the co-culture period (whose termination is defined hereafter as the time of fertilization), allowing us to perform a time-course observation of fertilization and embryogenesis (Figure 3D–3I).

**Figure 3.**
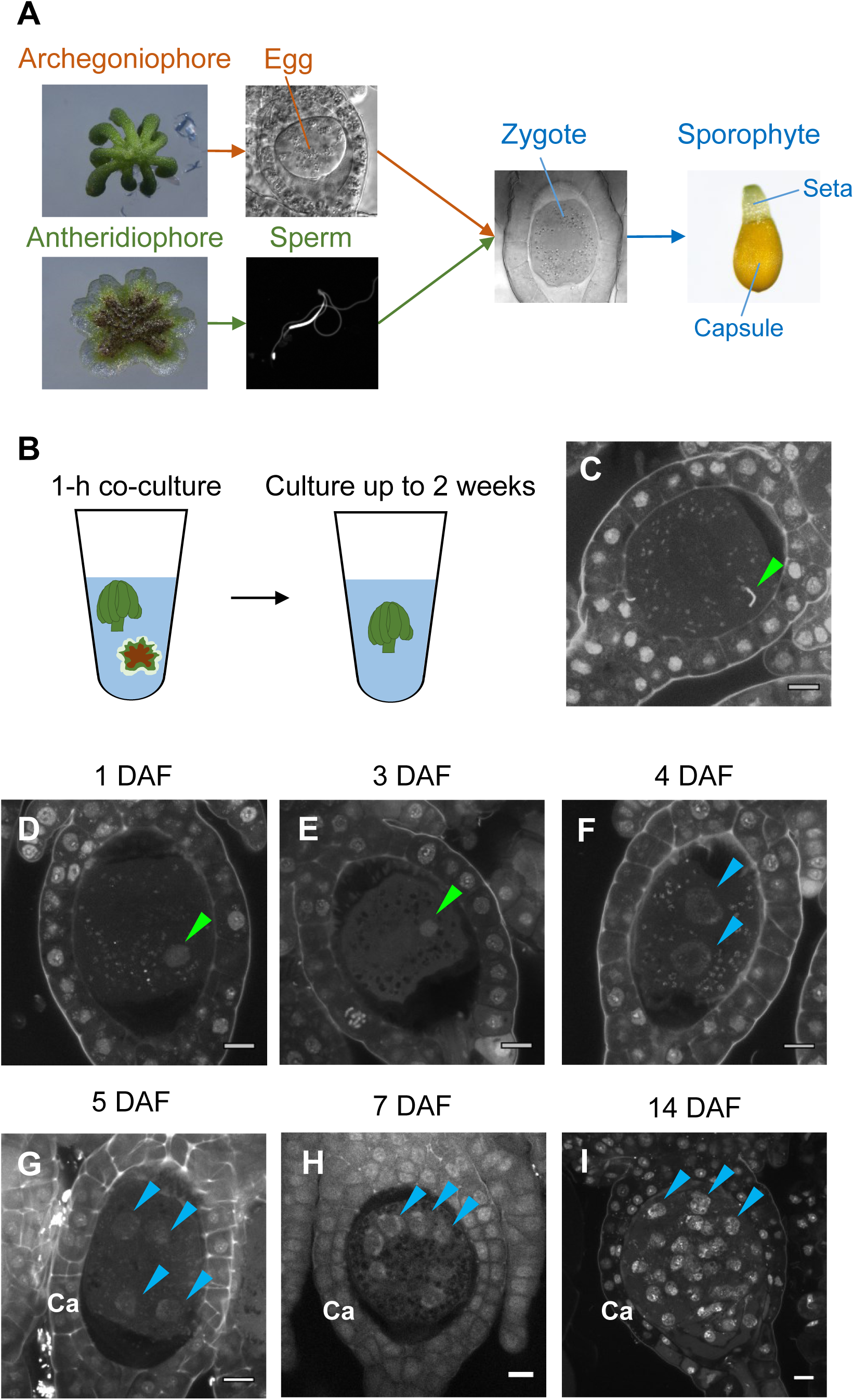
Time-course observation of subcellular dynamics during *M. polymorpha* fertilization. **A.** Schematic representation of sexual reproduction in *M. polymorpha*. Female plants develop umbrella-shaped sexual branches termed archegoniophores that form egg-containing archegonia. Male plants develop disc-shaped sexual branches termed antheridiophores that form antheridia, which produce numerous motile sperm cells. Upon soaking in water, the sperm cells are released from the antheridia and swim to egg cells in the archegonia. After fertilization, each zygote undergoes embryogenesis by dividing and differentiating into a sporophyte body consist of a capsule containing haploid spores and a short supportive stalk called the seta. **B.** Illustration of the in vitro fertilization method used in this study. Excised archegoniophores and antheridiophores were co-cultured in water for 1 h to allow fertilization to take place. The archegoniophores were transferred to a fresh tube containing water for further culturing. The tube lids were left open to allow gas exchange to occur. Archegoniophores containing sporophytes were cultured for up to 2 weeks. **C.** A DAPI-stained zygote after 1 h of co-culture. Most zygotes contained sperm nuclei at this time (green arrowhead). **D–I.** DAPI-stained zygotes and sporophytes at the indicated days after fertilization (DAF). Male pronuclei (green arrowheads) were visible at 1–3 DAF (D and E) in wild-type fertilized eggs. In most zygotes, karyogamy was completed, and cells were cleaved at 4 DAF (F). Sporophyte cells continued to divide at 5 to 14 DAF (G–I), as visualized by the presence of multiple nuclei (blue arrowheads; not all nuclei are labeled in H and I). Ca, calyptra. Bars, 10 µm.

We observed the cellular dynamics of zygotes and early embryos using optimized cell wall staining and tissue clearing techniques (Miyashima et al., 2019, Kurihara et al., 2015). No cell walls were stained in mature egg cells (Figure 3 - figure supplement 1A), indicating that cell walls are not present in mature eggs, a prerequisite for fusion with sperm cells. A cell wall around the zygote was detected only at 3 days after fertilization (DAF) (Figure 3 - figure supplement 1B–1D), suggesting that zygotic activation occurs slowly, requiring at least two days. The first zygotic division occurred at 4−5 DAF (Figure 3 - figure supplement 1E and 1F), a time point considerably later than that reported for the zygotes of flowering plants (20 hours after fertilization (HAF); Gooh et al., 2015, Uchiumi et al., 2007). At 1 to 3 DAF, a male pronucleus was clearly stained with DAPI, in contrast to the female pronucleus (not visible by DAPI staining), and was typically positioned halfway between the periphery and center of the fertilized egg (Figure 3D and 3E, green arrowhead). At 4 DAF, most zygotes completed the first division (Figure 3F), indicating that karyogamy takes place at 3–4 DAF. After 5 DAF, zygotes and the surrounding archegonial wall cells divided to form sporophytes and the calyptra (a protective gametophyte tissue), respectively (Figure 3G–3I). These cellular rearrangements, including karyogamy and embryogenesis, proceeded at a rate comparable to that in zygotes produced *in planta* (Figure 3 - figure supplement 2), confirming that our *in vitro* fertilization protocol faithfully recapitulated fertilization programs *in planta*.

### Maternal Mp*KNOX1* is required for pronuclear fusion in zygotes

To analyze the biological functions of Mp*KNOX1*, we generated loss-of-function mutants of Mp*KNOX1* using a CRISPR/Cas9 technique optimized for *M. polymorpha* (Sugano et al., 2018). We obtained two independent female mutant lines that harbored nucleotide insertions or deletions resulting in premature stop codons preceding the region encoding the HD (Figure 4 - figure supplement 1A). Both mutants were indistinguishable from wild-type females in terms of both vegetative and reproductive morphology (Figure 4 - figure supplement 2). Mature archegonia and egg cells of the Mp*knox1* mutants were also indistinguishable from those of the wild type (Figure 4A and 4D), indicating that Mp*KNOX1* functions are dispensable for both gametophyte development and gametogenesis.

**Figure 4.**
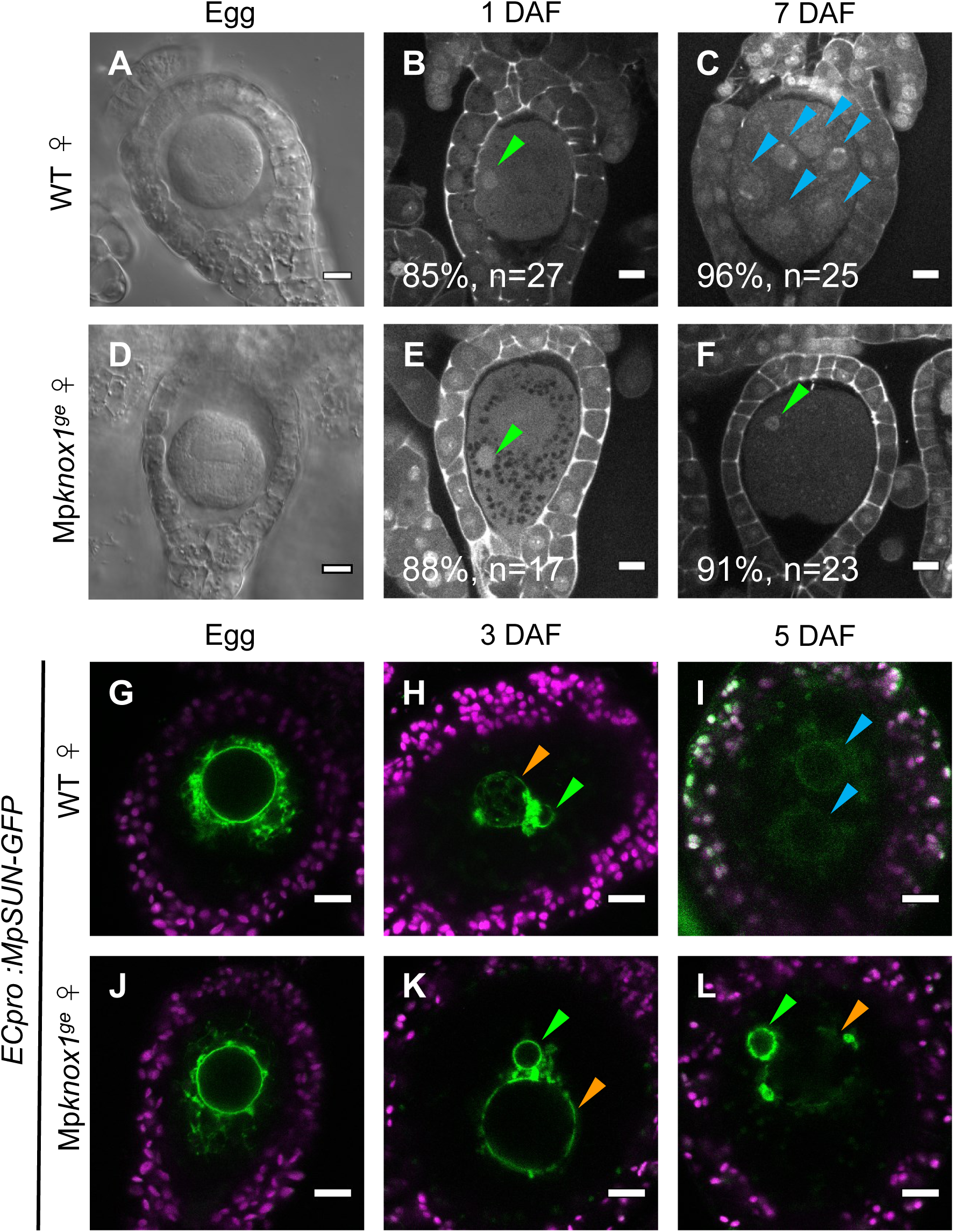
Maternally inherited Mp*KNOX1* is required for nuclear fusion. **A and D.** Bright-field images of wild-type (A) and Mp*knox1-1^ge^* (D) archegonia. **B, C, E, F.** 1-DAF (B, E) and 7-DAF (C, F) zygotes from a cross between wild-type female and male plants (B, C), and a cross between Mp*knox1-1^ge^* female and wild-type male plants (E, F), indicating that maternal Mp*KNOX1* is dispensable for fertilization (B, E) but is required for embryogenesis (C, F). **G–L.** Egg cells of *MpSUN-GFP* marker lines in the wild type (G) or Mp*knox1-2^ge^* (J) female background were crossed with wild-type males. At 3 DAF, male and female pronuclei were in contact with each other in both wild-type (H) and Mp*knox1* (K) eggs. At 5 DAF, zygotes derived from a wild-type egg started to divide (I), while those from an Mp*knox1* female (L) were arrested without nuclear membrane fusion. Green arrowhead, male pronucleus; orange arrowhead, female pronucleus; blue arrowhead, embryo nucleus. Bars, 10 µm.

We crossed the Mp*knox1* mutant females with wild-type males and observed the resulting zygotes by microcopy. Similar to wild-type zygotes, each Mp*knox1* egg fertilized with wild-type sperm harbored a male pronucleus at 1 DAF (85%, *n* = 27 for wild type and 88%, *n* = 17 for Mp*knox1*, Figure 4B and 4E), indicating that Mp*knox1* eggs are able to fuse with wild-type sperm and support decondensation of sperm nuclei. At 7 DAF, however, male and female pronuclei remained unfused in most fertilized Mp*knox1* eggs (91%, *n* = 23, Figure 4F), in contrast with wild-type fertilized eggs, which were undergoing sporophyte development (96%, *n* = 25) (Figure 4C). The two independent Mp*knox1* mutant lines exhibited indistinguishable defects in karyogamy. Importantly, this defect was rescued in archegonia expressing MpKNOX1-GFP driven by the Mp*KNOX1* promoter (*gMpKNOX1-GFP*, Figure 4 - figure supplement 3), confirming the notion that Mp*KNOX1* is required maternally to complete fertilization.

A small fraction of Mp*knox1* eggs that were fertilized with wild-type sperm developed into sporangia that produced functional spores (Figure 4 - figure supplement 4), suggesting that redundant genetic pathway(s) can compensate for the loss of Mp*KNOX1,* and/or that our Mp*knox1* mutant alleles were not null, though genomic sequences preceding the HD-coding region was disrupted. This slightly leaky penetrance allowed us to obtain male Mp*knox1* gametophytes from the resulting spores. The male Mp*knox1* mutants produced functional sperm capable of producing normal embryos when used to fertilize wild-type eggs (Figure 4 - figure supplement 5), indicating that paternal Mp*KNOX1* is dispensable for gametophyte development, fertilization, and embryogenesis. Together, these results indicate that the egg-derived functional Mp*KNOX1* allele or its protein products, but not those derived from sperm, are required to activate zygote development, more specifically karyogamy, during *M. polymorpha* fertilization.

In both animals and plants, karyogamy occurs via a two-step process: pronuclear migration and nuclear membrane fusion (Fatema et al., 2019). During pronuclear migration, one or both pronuclei migrate to become in close proximity, while during nuclear membrane fusion, the nuclear envelopes of the two pronuclei fuse together to produce a zygotic nucleus with both maternal and paternal genomes. To identify which of these steps is affected by the Mp*knox1* mutation, we visualized the pronuclear envelope by expressing GFP-tagged MpSUN proteins under the control of an egg-specific promoter (*ECpro:MpSUN-GFP*, see Materials and Methods for details). Before fertilization, wild-type and Mp*knox1* egg nuclei were of a similar size (approximately 20 µm in diameter) and were surrounded by a mesh-like membranous structure (Figure 4G and 4J). At 3 DAF, *ECpro:MpSUN-GFP* signals were visualized at the surfaces of both female and male pronuclei, suggesting the presence of a nuclear envelope (Figure 4H and 4K). In both wild-type- and Mp*knox1*-derived zygotes, female and male pronuclei were tethered to each other by a membranous structure marked by MpSUN-GFP (Figure 4H and 4K). At 5 DAF, however, male and female pronuclei remained separated by an intercalating membranous structure in Mp*knox1*-derived zygotes, while those from wild-type eggs already showed two embryonic nuclei (Figure 4I and 4L). Together, these observations indicate that maternal Mp*KNOX1* or its protein product is dispensable for the organization and migration of pronuclei but is required for pronuclear membrane fusion.

### Both maternal and paternal Mp*BELL* alleles contribute to karyogamy

KNOX proteins heterodimerize with BELL proteins to regulate gene transcription (Hay and Tsiantis, 2010). In *C. reinhardtii*, a *minus* gamete-derived KNOX protein (GSM1) heterodimerizes with a *plus* gamete-derived BELL protein (GSP1) upon fertilization and activates the majority of early zygote-specific genes (Joo et al., 2017, Kariyawasam et al., 2019, Lee et al., 2008, Lopez et al., 2015, Nishimura et al., 2012). Loss of GSM1 and/or GSP1 results in delayed pronuclear fusion, a phenotype similar to that of Mp*knox1* mutants. The phenotypic similarity between *M. polymorpha* Mp*knox1* and *C. reinhardtii gsm1* mutants suggests that the role of KNOX and BELL in zygote activation is conserved and that its evolutionary origin can be traced back to a common ancestor of the two species.

In support of this hypothesis, publicly available transcriptome data (Bowman et al., 2017) indicate that two of the five *BELL* genes of *M. polymorpha*, Mp*BELL3* (Mp8g02970) and Mp*BELL4* (Mp8g07680), are preferentially expressed in the antheridiophores of male plants, whereas Mp*BELL1* (Mp8g18310) and Mp*BELL5* (Mp5g11060) are preferentially expressed in sporophytes and archegonia, respectively (Figure 5 - figure supplement 1). RT-PCR analysis confirmed that Mp*BELL3* and Mp*BELL4* are specifically expressed in antheridiophores containing antheridia (Figure 5A).

**Figure 5.**
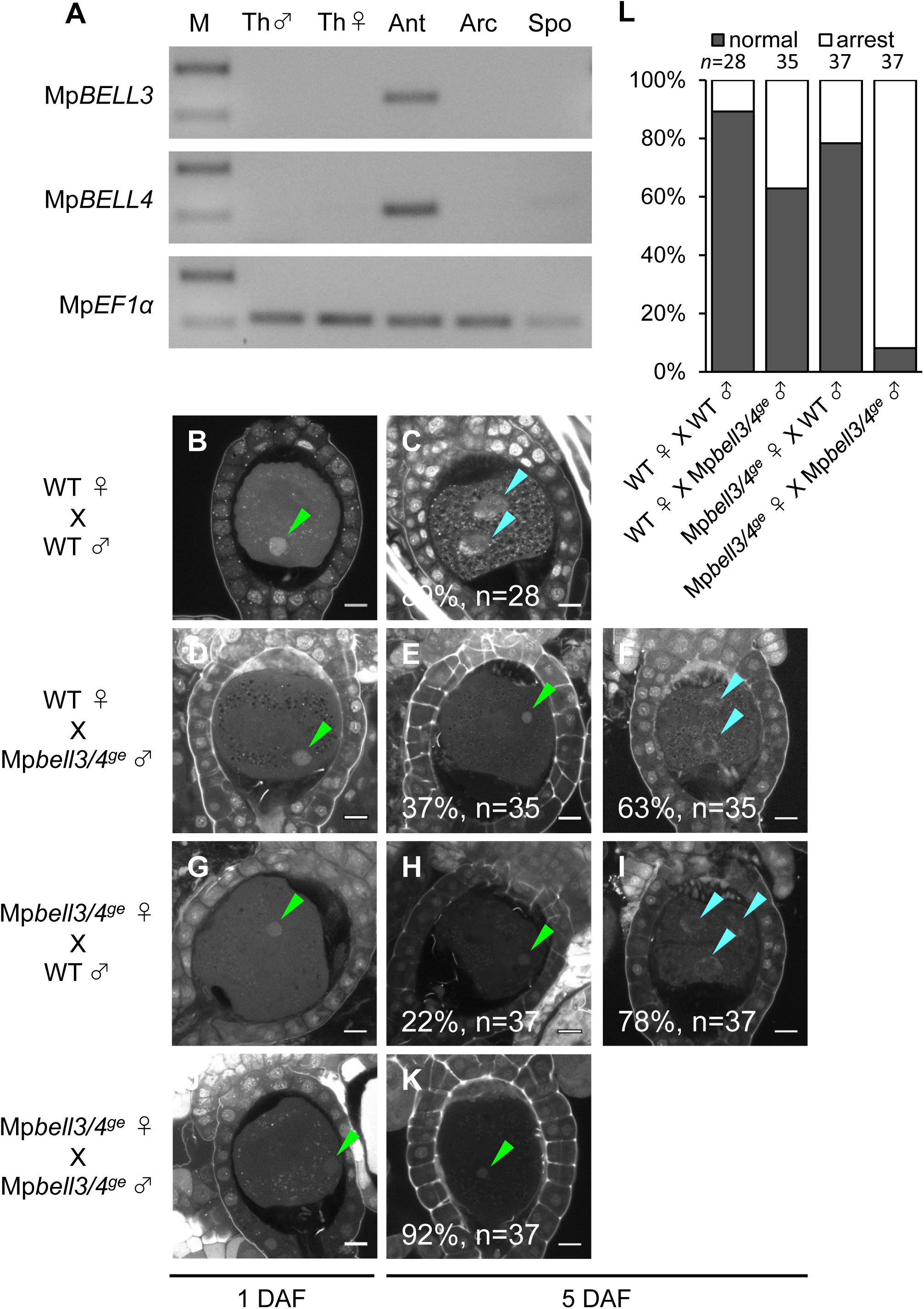
Both paternally and maternally inherited Mp*BELL* genes are required for karyogamy. **A.** RT-PCR analysis indicating that Mp*BELL3* and Mp*BELL4* are specifically expressed in antheridiophores. The lanes are labeled as in Figure 2A. Shown is a representative result from the experiments using three independently collected samples each with two technical replicates. **B–K.** Zygotes at 1 DAF (B, D, G, J) and 5 DAF (C, E, F, H, I, K) from crosses between a wild-type female and wild-type male (B, C) or Mp*bell3/4-2^ge^* male (D–F) and a Mp*bell3/4-2^ge^* female and a wild-type male (G–I) or Mp*bell3/4-2^ge^* male (J, K). The presence of male pronuclei (green arrowheads) in zygotes of all genotypes at 1 DAF (B, D, G, J) indicates that Mp*BELL3* and Mp*BELL4* are dispensable for plasmogamy. Note that zygotes produced from both or one Mp*bell3/4* parent exhibit a variable degree of karyogamy arrest, as visualized by the retention of male pronuclei (green arrowheads) among those starting embryonic division (nuclei labeled with blue arrowheads). Bars, 10 µm. **L.** Bar graph showing the ratios of zygotes with karyogamy arrest from the indicated crosses shown in B–K. Numbers of observed zygotes are shown above each bar.

To investigate whether Mp*BELL3* and/or Mp*BELL4* are required for fertilization, we generated two male and one female mutant lines in which Mp*BELL3* and Mp*BELL4* were simultaneously disrupted (hereafter referred to as Mp*bell3/4*) using a CRISPR/Cas9 nickase system (Figure 4 - figure supplement 1B and 1C). Both female and male Mp*bell3/4* gametophytes grew normally (Figure 4 - figure supplement 2) and produced normal gametangia and gametes capable of fertilization (Figure 5 -figure supplement 2, Figure 5B, D, G, and J). By contrast, approximately one-third of zygotes (37%, *n* = 35) produced by crossing wild-type females and Mp*bell3/4* males did not complete karyogamy by 5 DAF (Figure 5E, 5F and 5L).

Somewhat unexpectedly, zygotes obtained from a reciprocal cross (Mp*bell3/4* females with wild-type males) exhibited a low but significant degree of karyogamy arrest (22%, *n* = 37) (Figure 5H, 5I and 5L), despite the lack of detectable Mp*BELL3/4* expression in female gametes. Furthermore, a majority of 5-DAF zygotes obtained from a cross between Mp*bell3/4* females and Mp*bell3/4* males (92%, *n* = 37) were arrested at karyogamy (Figure 5K and 5L), as was the case for two independent male Mp*bell3/4* lines. These results suggest that not only paternally inherited Mp*BELL3* and/or Mp*BELL4* alleles(s) and/or their protein products, but also maternally inherited Mp*BELL3* and/or Mp*BELL4* allele(s) contribute to karyogamy.

### MpKNOX1 proteins transiently localize to male and female pronuclei in an Mp*BELL3/4*-dependent manner

Our genetic analyses indicated that Mp*KNOX1* functions after fertilization despite its specific transcription in unfertilized egg cells. This observation suggests that MpKNOX1 protein produced in unfertilized egg cells functions in zygotes. To test this hypothesis, we analyzed MpKNOX1 protein dynamics during fertilization by examining *gMpKNOX1-GFP* plants by confocal microscopy. In mature egg cells, MpKNOX1-GFP was specifically detected in the cytosol (Figure 6A). After crossing with a wild-type male, GFP signals were detected in both male and female pronuclei at 12 HAF (Figure 6B). At 24 HAF, the GFP signals were totally excluded from the pronuclei (Figure 6C). These observations suggest that upon fertilization MpKNOX1 translocates from the cytosol to pronuclei.

**Figure 6.**
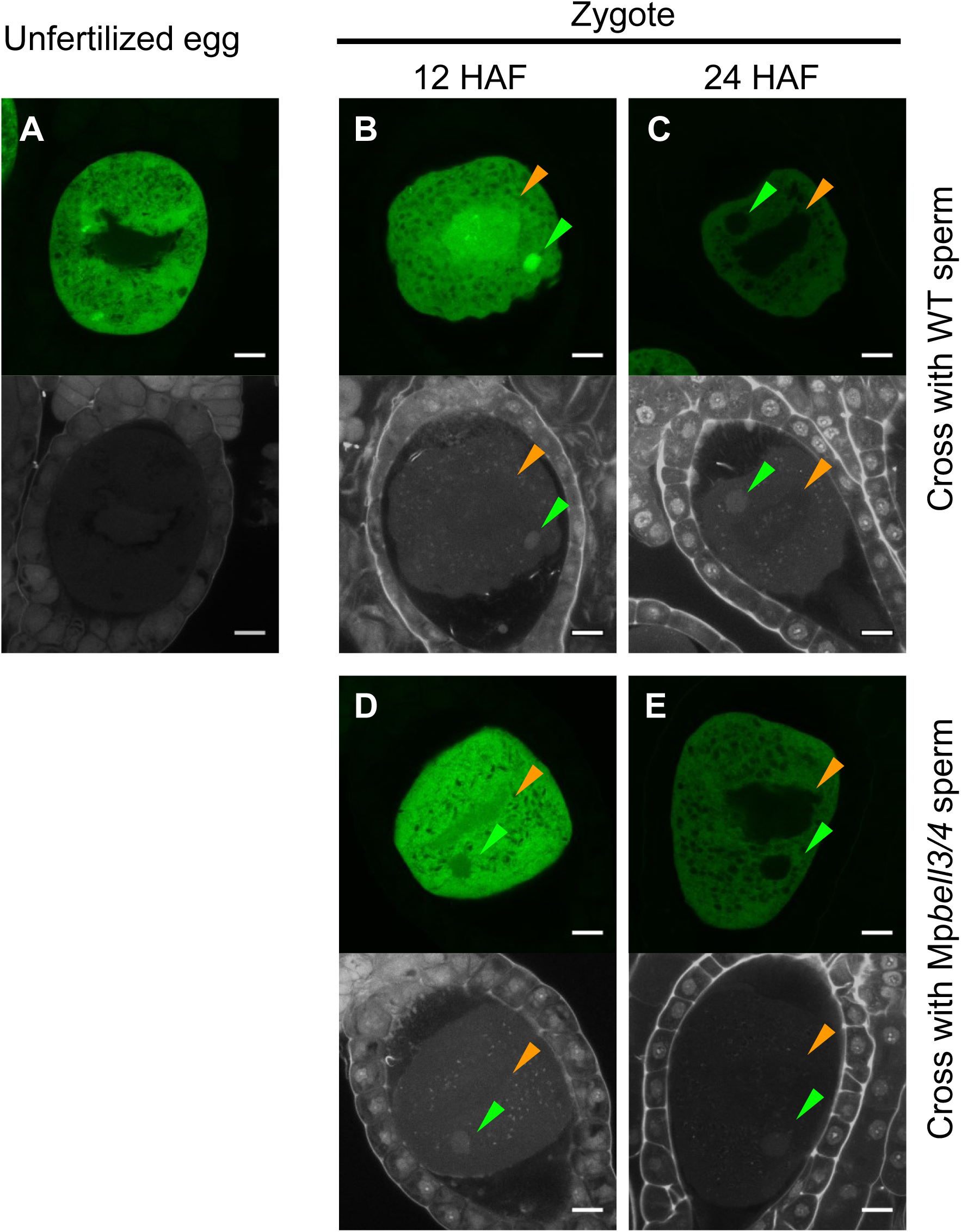
MpKNOX1 transiently localizes to female and male pronuclei prior to karyogamy. **A–E.** GFP (upper panels) and DAPI (lower panels) signals from *gMpKNOX1-GFP/*Mp*knox1-1^ge^* eggs (A) and zygotes obtained by crossing a *gMpKNOX1-GFP/*Mp*knox1-1^ge^* female with a wild-type (B, C) or Mp*bell3/4-2^ge^* (D, E) male. Note that before fertilization, MpKNOX1-GFP was exclusively localized to the cytosol (A). At 12 HAF, MpKNOX1-GFP signals were enriched in female (orange arrowhead) and male (green arrowhead) pronuclei in the wild-type background (B). In the absence of paternally inherited Mp*BELL3* and Mp*BELL4*, MpKNOX1-GFP remained mostly cytosolic (D), although weak GFP signals were detected in male and female pronuclei at 12 HAF (D). At 24 HAF, weak GFP signals were exclusively detected in the cytosol of both genotypes (C, E). Bars, 10 µm.

As KNOX proteins function as transcription factors, the transient localization of MpKNOX1 in pronuclei should be critical for its role in regulating karyogamy-promoting genes. In Arabidopsis, KNOX proteins are recruited to nuclei through interactions with BELL proteins. To examine whether paternally inherited Mp*BELL3* and/or Mp*BELL4* contribute to the pronuclear localization of MpKNOX1, we crossed *gMpKNOX1-GFP* females with wild-type or Mp*bell3/4* males and analyzed the subcellular localization of MpKNOX1-GFP in zygotes. In contrast to zygotes derived from a cross with wild-type males, where GFP signals were preferentially detected in male and female pronuclei, zygotes derived from a cross with Mp*bell3/4* males retained GFP signals in the cytosol (Figure 6B–6E). Together, these data suggest that paternally inherited Mp*BELL3* and/or Mp*BELL4* contribute to the pronuclear localization of MpKNOX1 after fertilization.

## Discussion

Here, we demonstrated that Mp*KNOX1* is an egg-specific gene in *M. polymorpha.* Mp*KNOX1* is strongly expressed in developing and mature eggs, whereas no expression was detected in gametophytes, sperm or sporophytes. The egg-specific expression of Mp*KNOX1* is in sharp contrast with the expression pattern of *KNOX1* genes in another model bryophyte, *P. patens*, where all three *KNOX1* genes are strongly expressed in sporophytes to regulate their development (Sakakibara et al., 2008). Rather, the egg-specific expression pattern of Mp*KNOX1* is reminiscent of that of *GSM1,* a *KNOX* gene of the unicellular green alga *C. reinhardtii* specifically expressed in *minus* gametes. Upon fertilization, GSM1 forms heterodimers with *plus* gamete-derived BELL protein GSP1 to activate expression of early zygote-specific genes (Joo et al., 2017, Lee et al., 2008, Nishimura et al., 2012). Accordingly, we suspected that MpKNOX1 might function as a gamete-derived zygote activator in *M. polymorpha*. Indeed, egg cells produced by two independent loss-of-function Mp*knox1* female mutants failed to produce embryos when fertilized with wild-type sperm. By contrast, wild-type eggs fertilized with Mp*knox1* sperm produced normal embryos and spores. These results clearly indicate that maternal Mp*knox1* exclusively contributes to embryogenesis.

The parent-of-origin effects of gene alleles can arise at several levels (Luo et al., 2014). In some cases, only one parental allele is transcribed in zygotes due to silencing of the other allele. In other cases, gene products such as proteins and small RNA molecules are synthesized in and/or carried over from gametes of a single sex. Our reporter analysis revealed that Mp*KNOX1* is preferentially transcribed in egg cells. By contrast, MpKNOX1-GFP accumulated in both unfertilized and fertilized eggs until 12 HAF, which mostly diminish at 24 HAF. These observations strongly argue for a mechanism in which MpKNOX1 produced in unfertilized eggs acts later in zygotes to initiate embryogenesis. This mode-of-action of MpKNOX1 is strikingly similar to that proposed for *C. reinhardtii* GSM1 (Lee et al., 2008).

How does egg-derived MpKNOX1 promote embryogenesis? We determined that in wild-type *M. polymorpha,* karyogamy is completed only after 3–4 DAF, a time considerably later than that reported for flowering plants (Gooh et al., 2015, Uchiumi et al., 2007). This slow progression of karyogamy is further delayed or arrested when maternal Mp*knox1* is mutated. Observation of nuclear membrane dynamics using an MpSUN-GFP marker revealed that in fertilized Mp*knox1* mutant eggs, karyogamy was arrested at the nuclear membrane fusion step, not the nuclear migration step. Consistent with the predicted role of MpKNOX1 as a transcription factor, MpKNOX1-GFP localized to both male and female pronuclei by 12 HAF, far before the initiation of nuclear membrane fusion. These observations suggest that gamete-derived MpKNOX1 functions in pronuclei to regulate the expression of gene(s) required for nuclear membrane fusion, thereby activating zygote development.

In both angiosperms and *C. reinhardtii*, the heterodimerization of KNOX and BELL is required for the nuclear translocation of these proteins, which in turn is required for them to regulate gene transcription (Hay and Tsiantis, 2010, Lee et al., 2008). Consistent with this notion, our genetic and imaging analyses suggested that sperm-derived MpBELLs are required to recruit MpKNOX1 to male and female pronuclei. Among the five *BELL* genes in the *M. polymorpha* genome, Mp*BELL3* and Mp*BELL4* are specifically expressed in male sexual organs. Mp*bell3/4* mutant males could produce motile sperm, but approximately 40% of wild-type eggs fertilized with Mp*bell3/4* sperm did not produce embryos compared to ∼10% of those fertilized with wild-type sperm. This partial loss of embryogenic ability was accompanied by severely compromised pronuclear localization of MpKNOX1-GFP. These observations strongly suggest a mechanism in which paternal MpBELL3 and/or MpBELL4 recruit maternal MpKNOX1 to pronuclei.

Interestingly, however, our genetic analyses also indicated that not only paternal but also maternal Mp*BELL3* and/or Mp*BELL4* are required for karyogamy with full penetrance, even though their expression was not detected in maternal organs or egg cells. A plausible explanation for this observation is that maternally inherited Mp*BELL3* and/or Mp*BELL4* alleles become transcriptionally activated in zygotes and that Mp*BELL3* and/or Mp*BELL4* expression post-fertilization acts to replenish MpBELL proteins to ensure karyogamy with high penetrance.

Based on these observations, we propose a two-step model of MpBELL3/4 activity (Figure 7A). According to this model, sperm-borne MpBELL3 and MpBELL4 proteins heterodimerize with egg-derived MpKNOX1 upon fertilization and activate the transcription of zygote-specific genes as well as maternal Mp*BELL3* and/or Mp*BELL4.* MpBELL3 and MpBELL4 produced *de novo* further activate the expression of zygote-specific genes, including genes required for karyogamy. Such feed-forward regulation would efficiently compensate for the presumably small amounts of MpBELL proteins inherited from the sperm cytosol. Considering that a recent phenotypic analysis of newly isolated *gsm1* and *gsp1* alleles revealed the biparental contribution of GSM1 to karyogamy in *C. reinhardtii* (Kariyawasam et al., 2019), our findings further emphasize the similarity of KNOX/BELL-mediated zygote activation between *M. polymorpha* and *C. reinhardtii*.

**Figure 7.**
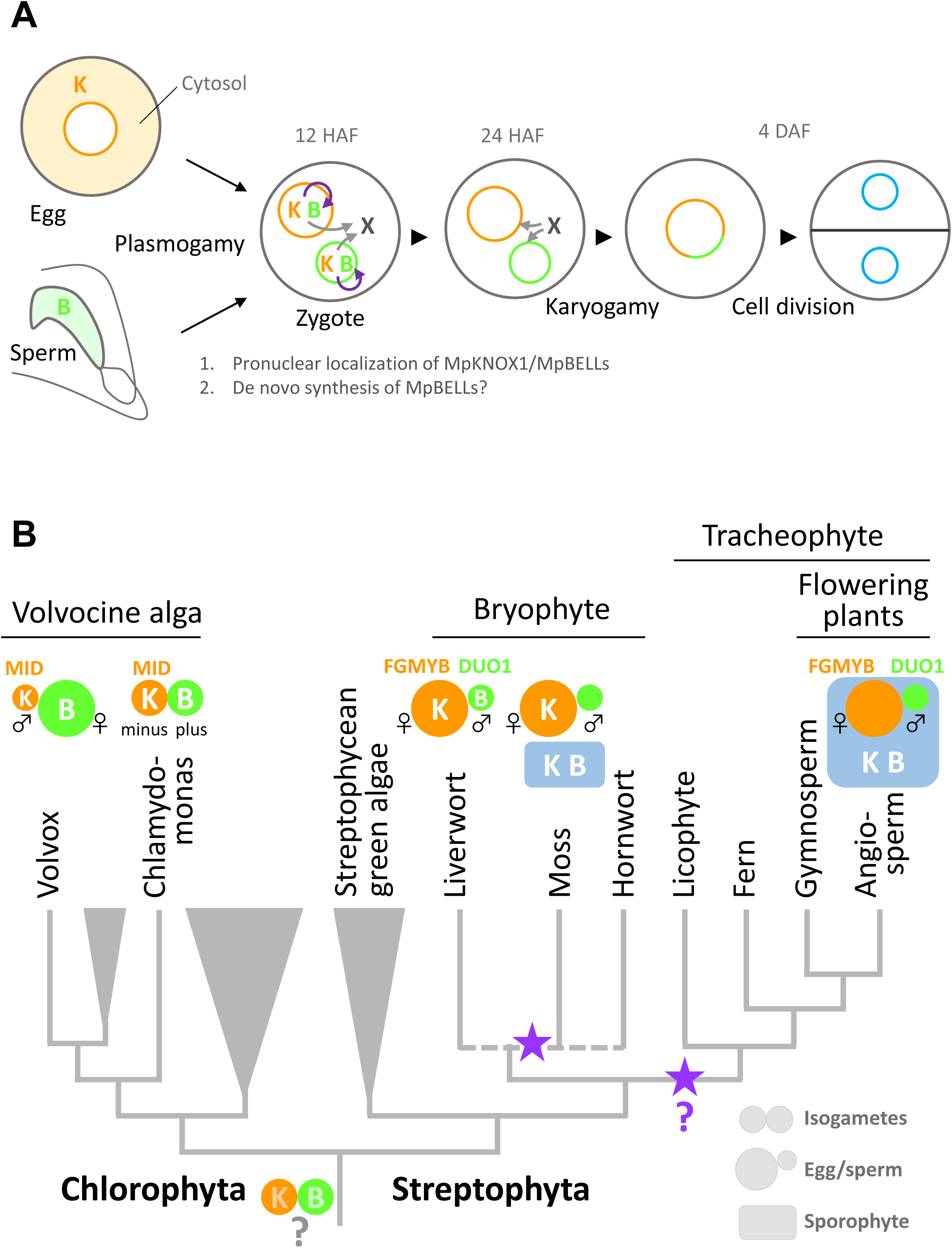
Functions and expression patterns of KNOX/BELL transcription factors in green plants. **A.** Expression patterns of KNOX and BELL proteins and their predicted role in zygote activation in *M. polymorpha.* “K” and “B” represent MpKNOX1 and MpBELL protein subunits, respectively. Orange, green and blue circles represent female pronuclei, male pronuclei and nuclei of embryo cells, respectively. Purple curved arrows indicate auto-amplification of BELL levels by KNOX/BELL-mediated transcriptional control. X indicates unknown karyogamy-promoting factor(s) whose expression and/or functions are activated by KNOX/BELL-mediated transcription. **B.** Predicted evolutionary trajectory of KNOX/BELL expression patterns along the green plant lineages. Orange/green circles and blue rectangles represent gametes and sporophyte bodies, respectively. “K” and “B” indicate the expression of KNOX and BELL proteins, respectively. Purple stars indicate the predicted positions at which the functional transition of KNOX/BELL from zygote activation to sporophyte morphogenesis occurred. Also indicated are the expression patterns of evolutionarily conserved regulators of sexual differentiation; a male-determinant factor MID of volvocine algae, and female- and male-differentiation factors, FGMYB and DUO1, respectively, of land plants. Note that the KNOX/BELL expression patterns in ancestral plants at the bottom of the tree is an inference.

While our study revealed the striking conservation of KNOX/BELL functions between *M. polymorpha* and *C. reinhardtii,* this finding is somewhat unexpected from a phylogenetic viewpoint, because KNOX/BELL proteins in the model bryophyte *P. patens* control sporophyte development, as do KNOX/BELL proteins in angiosperms. Outside the plant kingdom, TALE TFs in yeasts and fungi promote the haploid-to-diploid transition (Goutte and Johnson, 1988, Herskowitz, 1989, Kües et al., 1992, Spit et al., 1998). Thus, perhaps the shared zygote-activating functions of KNOX/BELL in *M. polymorpha* and *C. reinhardtii* reflect an ancestral state (Bowman et al., 2016).

Considering the generally accepted view of bryophyte monophyly (de Sousa et al., 2019, Puttick et al., 2018), our findings suggest that the functional transition of KNOX/BELLs from zygote activation to sporophyte morphogenesis occurred at least twice during land plant evolution, including once in the bryophyte lineage and once in the tracheophyte lineage (Figure 7B). Among the three bryophyte clades, liverworts have simpler sporophyte morphology than mosses and hornworts (Shimamura, 2016). In mosses, the extension of sporangia from gametophore apices depends on the activity of the seta meristem, where KNOX1 proteins play a role in promoting cell division, whereas the seta of *M. polymorpha* elongates exclusively by cell elongation (Ligrone et al., 2012b, Ligrone et al., 2012a, Sakakibara et al., 2008). Thus, the evolutionary innovation of meristematic tissues might have promoted the functional transition of KNOX/BELL from zygote activation to sporophyte morphogenesis.

Our study revealed that the zygote-activating function of KNOX/BELL is conserved between *C. reinhardtii* and *M. polymorpha*, which belong to Chlorophyta and Streptophyta, respectively. These two major green plant lineages were separated more than 700 million years ago (Becker, 2013) (Figure 7B). Interestingly, however, the sex-specific expression patterns of KNOX and BELL in *C. reinhardtii* are opposite to those in *M. polymorpha*. In *C. reinhardtii*, KNOX (GSM1) and BELL (GSP1) are expressed in isogamous *minus* and *plus* gametes (Lee et al., 2008), which directly evolved into male and female gametes, respectively, in oogamous (with small motile gametes and large immotile gametes) *Volvox carteri,* primarily by modifying genes acting downstream of the conserved sex-determinant protein MID (Ferris and Goodenough, 1997, Geng et al., 2014, Geng et al., 2018, Nozaki et al., 2006) (Figure 7B). Results from other research groups indicate that the expression specificity of KNOX/BELL is conserved along the *volvocine* lineage (with *minus* and male gametes expressing GSM1; personal communication with Takashi Hamaji (Kyoto University, Japan) and James Umen (Donald Danforth Plant Science Center, MO)). Thus, at least in one Chlorophyta lineage, KNOX and BELL are expressed in male and female gametes, respectively, a situation opposite to that in *M. polymorpha (KNOX* for females and *BELL* for males) (Figure 7B).

In land plants, oogamy likely evolved once, as key regulators of gametophytic sexual differentiation, such as FGMYBs for females and DUO POLLEN 1 (DUO1) for males, are shared between *M. polymorpha* and Arabidopsis (see Hisanaga et al., 2019b for a review) (Figure 7B). Theoretical analyses predicted that anisogamy evolved through disruptive selection, where an increased volume of one type of gamete favors zygote fitness, allowing the other gamete type to increase in number at the expense of volume (Parker et al., 1972). Thus, in ancestral green plants, KNOX/BELL expression specificity did not affect gamete morphology or function. This notion is consistent with our observation that neither Mp*KNOX1* nor Mp*BELL3/4* contribute to gamete development or function in extant *M. polymorpha*.

In summary, our study revealed a critical role of *KNOX* and *BELL* in zygote activation in the model bryophyte *M. polymorpha,* an early-diverging land plant. This is in stark contrast with a well-recognized role of *KNOX*/*BELL* in sporophytic meristem maintenance in angiosperms and another model bryophyte *P. patens*. Rather, striking conservation of KNOX/BELL functions in the promotion of karyogamy across phylogenetically distant *M. polymorpha* and *C. reinhardtii* suggests that functions of KNOX/BELL heterodimers shifted from zygote-activation to sporophyte development as land plant evolved. This view is consistent with a proposed evolutionary scenario of HD proteins in a broader eukaryotic taxa, including fungi and metazoans (Bowman et al., 2016).

## Materials and methods

### Plant materials

*Marchantia polymorpha* L. subsp. *ruderalis* accessions Takaragaike 1 (Tak-1) and Takaragaike 2 (Tak-2; (Ishizaki et al., 2016)) were used as the wild-type male and female, respectively. Plants were cultured on half-strength Gamborg’s B5 medium solidified with 1% (w/v) agar under continuous white light at 22 °C. To induce reproductive development, 10-day-old thalli were transferred to a pot containing vermiculite and grown under white light supplemented with far-red illumination generated by LED (VBL-TFL600-IR730; IPROS Co., Tokyo, Japan).

### Archegonium sampling, RNA extraction and Illumina sequencing

Mature archegonia were manually dissected from wild-type archegoniophores and divided into four pools, each composed of approximately 1,000 archegonia. Mature archegonia of Mp*rkd-1* and Mp*rkd-3* (Koi et al., 2016) were collected and each divided into two pools of approximately 1,000 archegonia. Total RNA was extracted from the samples with an RNeasy Plant Mini Kit (Qiagen, Venlo, Netherlands) according to the manufacturer’s protocol. The quality and quantity of the RNA were evaluated using a Qubit 2.0 Fluorometer (Life Technologies, Carlsbad, CA) and a RNA6000 Nano Kit (Agilent Technologies, Santa Clara, CA). Sequence libraries were constructed with a TruSeq RNA Sample Prep Kit v2 (Illumina, San Diego, CA) according to the manufacturer’s protocol. The quality of each library was examined using a Bioanalyzer with High Sensitivity DNA Kit (Agilent Technologies) and a KAPA Library Quantification Kit for Illumina (Roche diagnostics, Basel, Switzerland). An equal amount of each library was mixed to generate a 2-nM pooled library. Next-generation sequencing was performed using the HiSeq 1500 platform (Illumina, San Diego, CA) to generate 126-nt single-end data. Sequence data have been deposited at the DDBJ BioProject and BioSample databases under accession numbers PRJDB9329 and SAMD00205647-SAMD00205654, respectively.

### Data analysis

Read data were mapped to the genome sequence of *M. polymorpha* v3.1 using TopHat ver. 2.0.14 (Trapnell et al., 2009) with default parameters. FPKM values were calculated using the DESeq2 package in R (Love et al., 2014). Differentially expressed genes (DEGs) were identified using the TCC package in R (Sun et al., 2013) with a criterion of FDR < 0.01. Candidate egg-specific genes were identified by filtering the DEGs with a threshold of Mp*rkd*/WT ratio < −3 and FPKM in WT > 1.

### Semi in vitro culture and genetic crossing

Mature archegoniophores and antheridiophores were separated from thalli and collected into a 5-mL plastic tube containing 3 mL of water. Following co-culture for 1 h under white light at 22 °C, the archegoniophores were transferred to new 5 mL plastic tubes containing 3 mL of water and cultured under white light at 22 °C prior to observation.

### Microscopy

To observe MpSUN-GFP expression, archegonia were excised under a dissecting microscope and mounted in half-strength Gamborg’s B5 liquid medium. To observe pronuclei, archegoniophores were dissected and soaked in PFA fixative solution (4% [w/v] paraformaldehyde, 0.1% [w/v] DAPI, and 0.01% [v/v] Triton X-100 in 1 x PBS buffer). The samples were briefly vacuum-infiltrated four times and incubated for 1 h at room temperature with gentle shaking. The samples were washed twice with 1 x PBS and cleared by incubating in ClearSee solution containing 0.1% (w/v) DAPI for 2–3 days (Kurihara et al., 2015). The cleared samples were observed under a Nikon C2 confocal laser-scanning microscope (Nikon Instech, Tokyo, Japan). Sperm cells were stained with DAPI as described previously (Hisanaga et al., 2019a).

### RT-PCR

RNA extraction, cDNA synthesis and RT-PCR were performed as described previously (Hisanaga et al., 2019a) using the primer sets listed in Table S2.

### DNA constructs

The plasmids used in this study were constructed using the Gateway^TM^ cloning system (Ishizaki et al., 2015), the SLiCE method (Motohashi, 2015) or Gibson assembly (Gibson et al., 2009). The primers used for DNA construction are listed in Table S2.

#### pMpGE010_MpKNOX1ge

A DNA fragment producing Mp*KNOX1*-tageting gRNAs was prepared by annealing a pair of synthetic oligonucleotides (MpKNOX1ge01Fw/MpKNOX1ge01Rv). The fragment was inserted into the BsaI site of pMpGE_En03 (Cat. No. 71535, Addgene, Cambridge, MA) to yield pMpGE_En03-MpKNOX1ge, which was transferred into pMpGE010 (Cat. No. 71536, Addgene) (Sugano et al., 2018) using the Gateway^TM^ LR reaction (Thermo Fisher Scientific, Waltham, MA) to generate pMpGE010_MpKNOX1ge.

#### pMpGE017_MpBELL4ge-MpBELL3ge

To construct a plasmid to disrupt Mp*BELL3* and Mp*BELL4* simultaneously, four oligonucleotide pairs (MpBELL4-ge1-Fw/MpBELL4-ge1-Rv, MpBELL4-ge2-Fw/MpBELL4-ge2-Rv, MpBELL3-ge3-Fw/MpBELL3-ge3-Rv and MpBELL3-ge4-Fw/MpBELL3-ge4-Rv) were annealed and cloned into pMpGE_En04, pBC-GE12, pBC-GE23 and pBC-GE34 to yield pMpGE_En04-MpBELL4-ge1, pBC-GE12-MpBELL4-ge-2, pBC-GE23-MpBELL3-ge3 and pBC-GE34-MpBELL3-ge4, respectively. These four plasmids were assembled via BglI restriction sites and ligated to yield pMpGE_En04-MpBELL4-ge12-MpBELL3-ge34. The resulting DNA fragment containing four Mp*U6*promoter-gRNA cassettes was transferred into pMpGE017 using the Gateway^TM^ LR reaction to yield pMpGE017_MpBELL4-ge12-MpBELL3-ge34.

#### pMpSL30_MpKNOX1pro-H2B-GFP-3’MpKNOX1

A 6.3-kb genomic fragment spanning the 5.3-kb 5′ upstream sequence plus the 1-kb 5′-UTR of Mp*KNOX1* was amplified from Tak-2 genomic DNA using the primers H-MpKNOX1pro-Fw and SmaI-MpKNOX1pro-Rv. A vector backbone containing the GFP-coding sequence was prepared by digesting pAN19_SphI-35S-lox-IN-lox-NosT-SmaI-GFP-NaeI (a kind gift from Dr. Shunsuke Miyashima) with SphI and SmaI. The two fragments were assembled using the SLiCE reaction to yield pAN19_MpKNOX1pro-SmaI-GFP-NaeI. The Histone H2B-coding sequence from Arabidopsis was amplified from the pBIN41_DUO1pro-H2B-YFP-nos vector (Hisanaga, unpublished) using the primers KNOXp-H2B-Fw and GFP-H2B-Rv. The fragment was inserted into the SmaI site of pAN19_MpKNOX1pro-SmaI-GFP-NaeI by the SLiCE reaction, yielding pAN19_MpKNOX1pro-H2B-GFP-NaeI. A 4-kb fragment containing the 0.5-kb 3′-UTR plus 3.5-kb 3′-flanking sequences of Mp*KNOX1* was amplified from Tak-2 genomic DNA using the primers G-MpKNOX1ter-Fw and E-MpKNOX1ter-Rv. The fragment was inserted into the NaeI site of pAN19_MpKNOX1pro-H2B-GFP-NaeI by the SLiCE reaction, yielding pAN19_MpKNOX1pro-H2B-GFP-3’MpKNOX1. The MpKNOX1pro-H2B-GFP-3’MpKNOX1 fragment was excised from pAN19_MpKNOX1pro-H2B-GFP-3’MpKNOX1 by digestion with AscI and inserted into pMpSL30 (Hisanaga et al., 2019a) to yield pMpSL30_MpKNOX1pro-H2B-GFP-3’MpKNOX1.

#### gMpKNOX1-GFP

A 3.4-kb genomic fragment spanning the entire exon and intron region of Mp*KNOX1* was amplified from Tak-2 genomic DNA using the primers gMpKNOX1-Fw and gMpKNOX1-Rv. The fragment was inserted into the SmaI site of pAN19_MpKNOX1pro-SmaI-GFP-NaeI by the SLiCE reaction to yield pAN19_MpKNOX1pro-MpKNOX1-GFP-NaeI. A 4-kb fragment containing the 0.5-kb 3′-UTR and 3.5-kb 3′-flanking sequences of Mp*KNOX1* was amplified from Tak-2 genomic DNA using the primers G-MpKNOX1ter-Fw and E-MpKNOX1ter-Rv. The fragment was inserted into the NaeI site of pAN19_MpKNOX1pro-MpKNOX1-GFP-NaeI by the SLiCE reaction, yielding pAN19_gMpKNOX1-GFP. The gMpKNOX1-GFP fragment was excised from pAN19_gMpKNOX1-GFP by digestion with AscI and inserted into pMpSL30 to yield pMpSL30_gMpKNOX1-GFP.

#### ECpro:MpSUN-GFP

A 5-kb genomic fragment spanning the 3.4-kb 5′ upstream and 1.6-kb 5′-UTR sequences of Mp5g18000 was amplified from Tak-2 genomic DNA using the primers H-ECpro-Fw and G-SpeI-ECpro-Rv. The fragment was inserted into the SpeI site of pAN19_SpeI-GFP-NaeI by the SLiCE reaction, yielding pAN19_ECpro-SpeI-GFP-NaeI. The 1.5-kb Mp*SUN* (Mp5g02400)-coding sequence was amplified from a cDNA library of *M. polymorpha* using the primers E-MpSUN-Fw and G-MpSUN-Rv. The fragment was inserted into the SpeI site of pAN19_ECpro-SpeI-GFP-NaeI by the SLiCE reaction, yielding pAN19_ECpro-MpSUN-GFP. The ECpro-MpSUN-GFP fragment was excised from pAN19_ECpro-MpSUN-GFP by digestion with AscI and inserted into pMpSL30 to yield pMpSL30_ ECpro-MpSUN-GFP.

### Generation of transgenic *M. polymorpha*

Genome editing constructs were introduced into *M. polymorpha* sporelings as described previously (Ishizaki et al., 2008). Other constructs were introduced into regenerating thalli (Kubota et al., 2013) or gemmae using the G-AgarTrap method (Tsuboyama et al., 2018).

## Acknowledgments

We thank Masako Kanda for technical assistance and Shunsuke Miyashima for DNA materials. This work was supported by MEXT KAKENHI grants 17J08430 to T.H., 25113007 to K.N., 17H05841 and 18K06285 to S.Y., and 25113009 and 17H07424 to T.K. T.H. was supported by a JSPS Fellowship for Young Scientists and a funding from the European Union’s Framework Programme for Research and Innovation Horizon 2020 (2014-2020) under the Marie Curie Skłodowska Grant Agreement Nr. 847548.

## Author contributions

T.H., S.F., Y.C., K.S., and S.Y. performed the experiments. R.S. analyzed RNA-seq data. T.H., T.K., F.B., and K.N. designed the project. T.H. and K.N. wrote the manuscript. All the authors jointly interpreted the data and thoroughly checked the manuscript.

## Conflict of interest

Authors declare no conflict of interests.

---

**Figure 1 - figure supplement 1.**
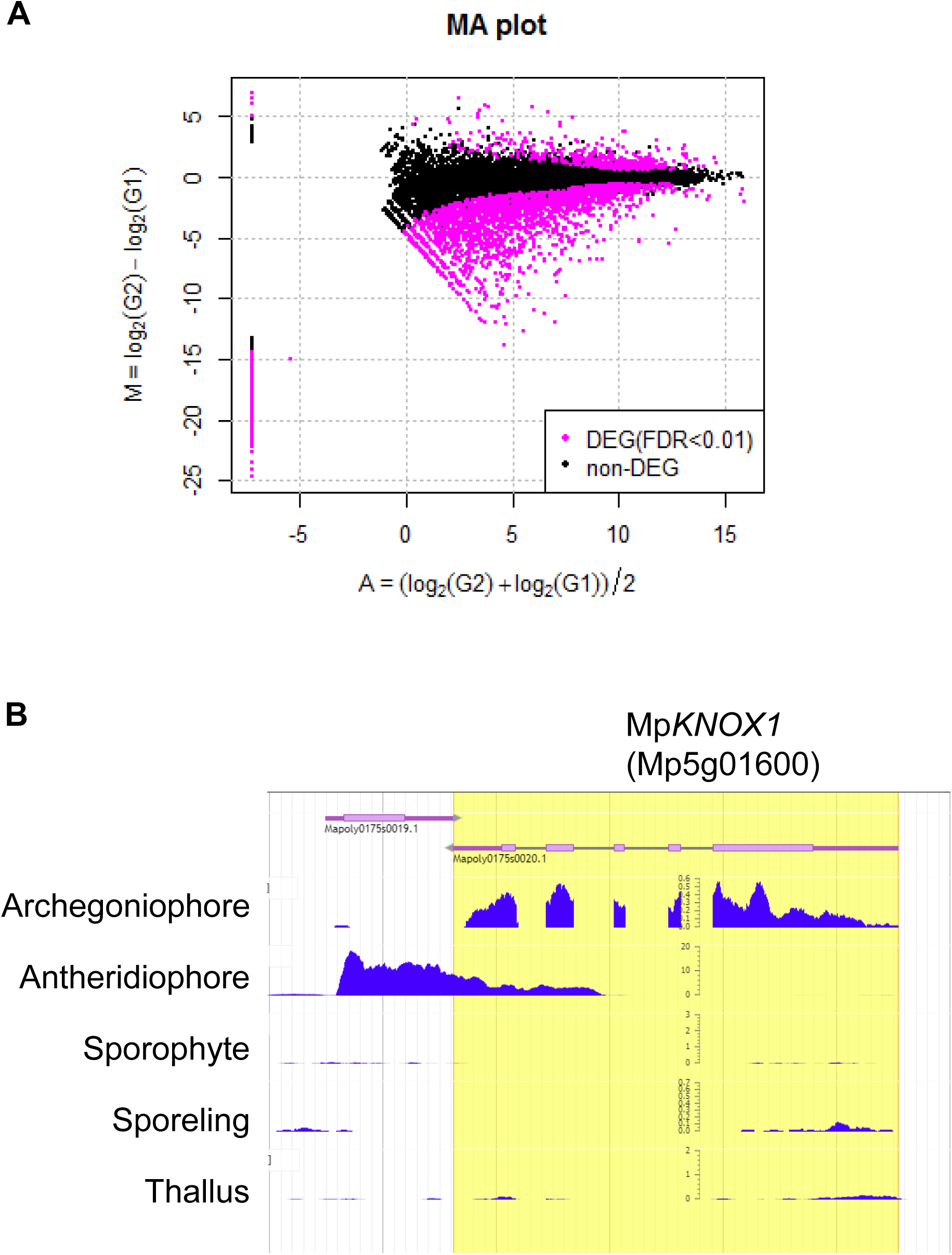
Comparative transcriptome analysis of wild-type and Mp*rkd* archegonia and identification of Mp*KNOX1* as an egg-specific gene in *M. polymorpha*. **A.** M-A plot of differential expression analysis between wild-type and Mp*rkd* archegonia. Differentially expressed genes (DEGs) are plotted in magenta. **B.** Captured Genome Browser image showing the expression levels along the Mp*KNOX1* locus in the indicated tissue types (archegoniophore; SRX301555, antheridiophore; SRX301553, sporophyte; SRX301556, sporeling; SRX301559, thallus; SRX301557). Mp*KNOX1* is specifically expressed in archegoniophores.

**Figure 3 - figure supplement 1.**
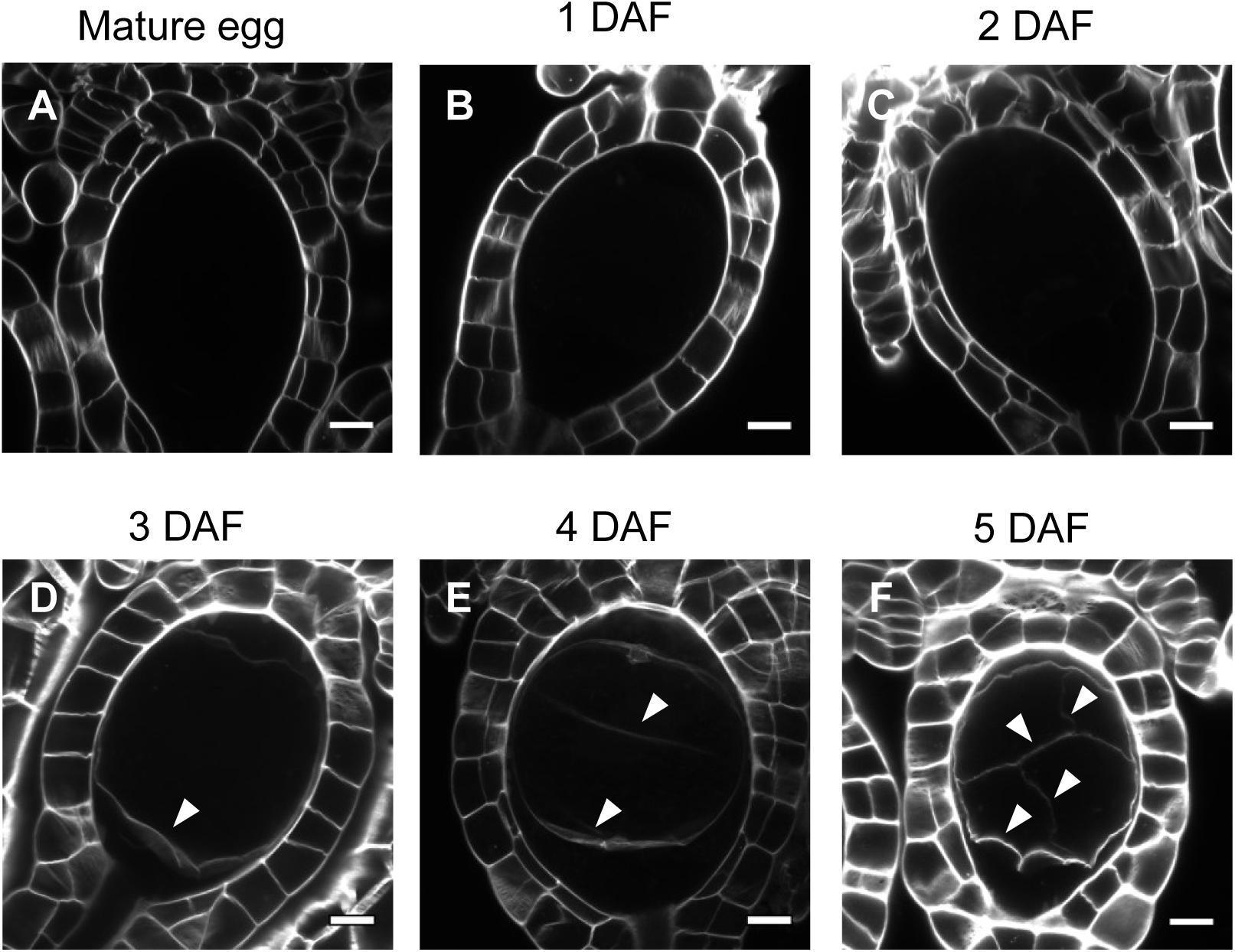
Cell wall regeneration during zygote development. Confocal images of archegonia hosting an egg cell, zygotes or embryos. Cell walls stained with SCRI Renaissance 2200 were detected only after embryogenesis (arrowheads), but not in mature eggs or zygotes at 3 DAF or earlier. Bars, 10 µm.

**Figure 3 - figure supplement 2.**
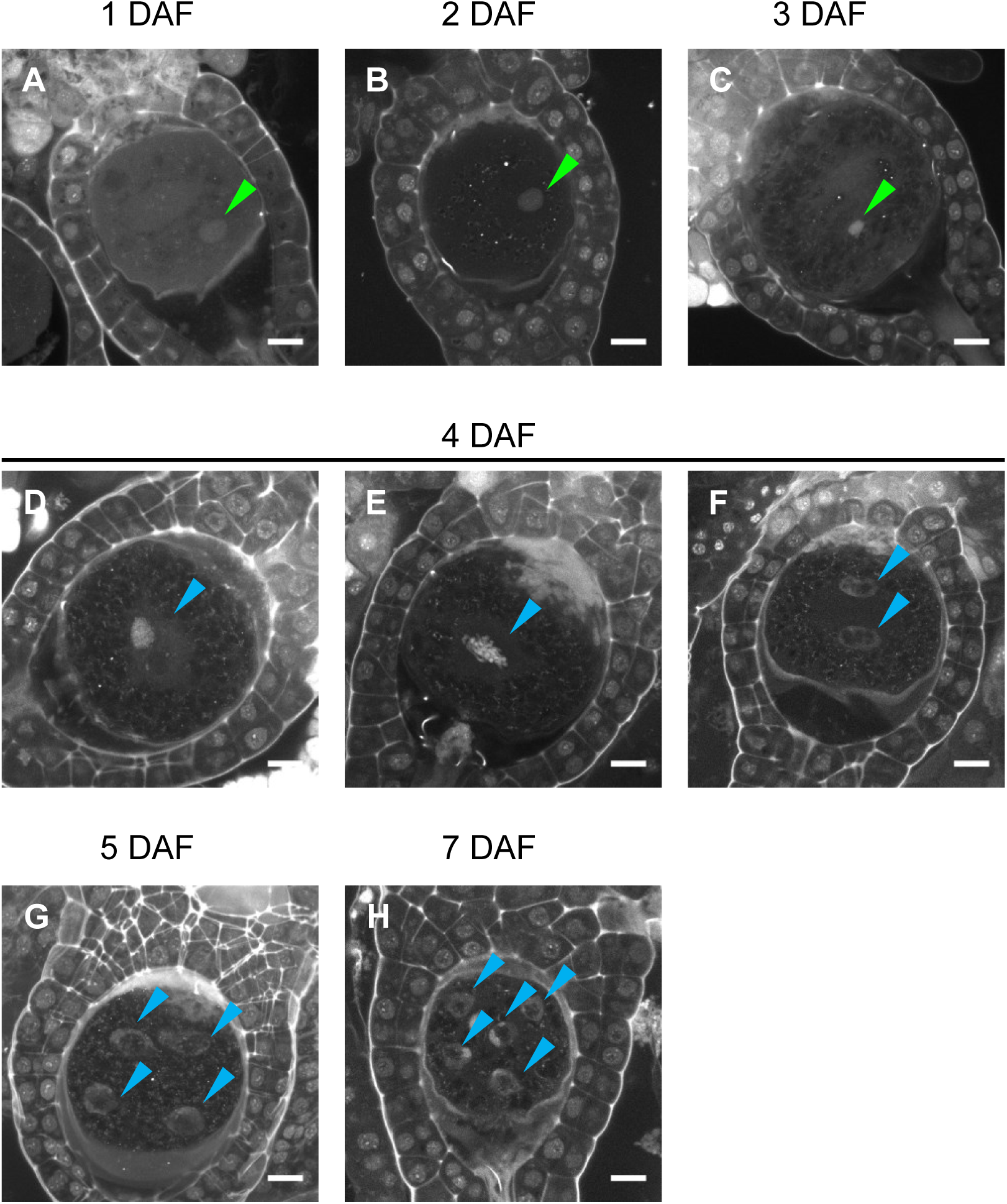
Cellular dynamics of zygotes and embryos generated by *in planta* crossing. DAPI-stained zygotes and embryos derived from in planta crosses at the indicated stages, showing that the timing of subcellular dynamics was equivalent to that generated by the in vitro fertilization method shown in Figure 3. Bars, 10 µm.

**Figure 4 - figure supplement 1.**
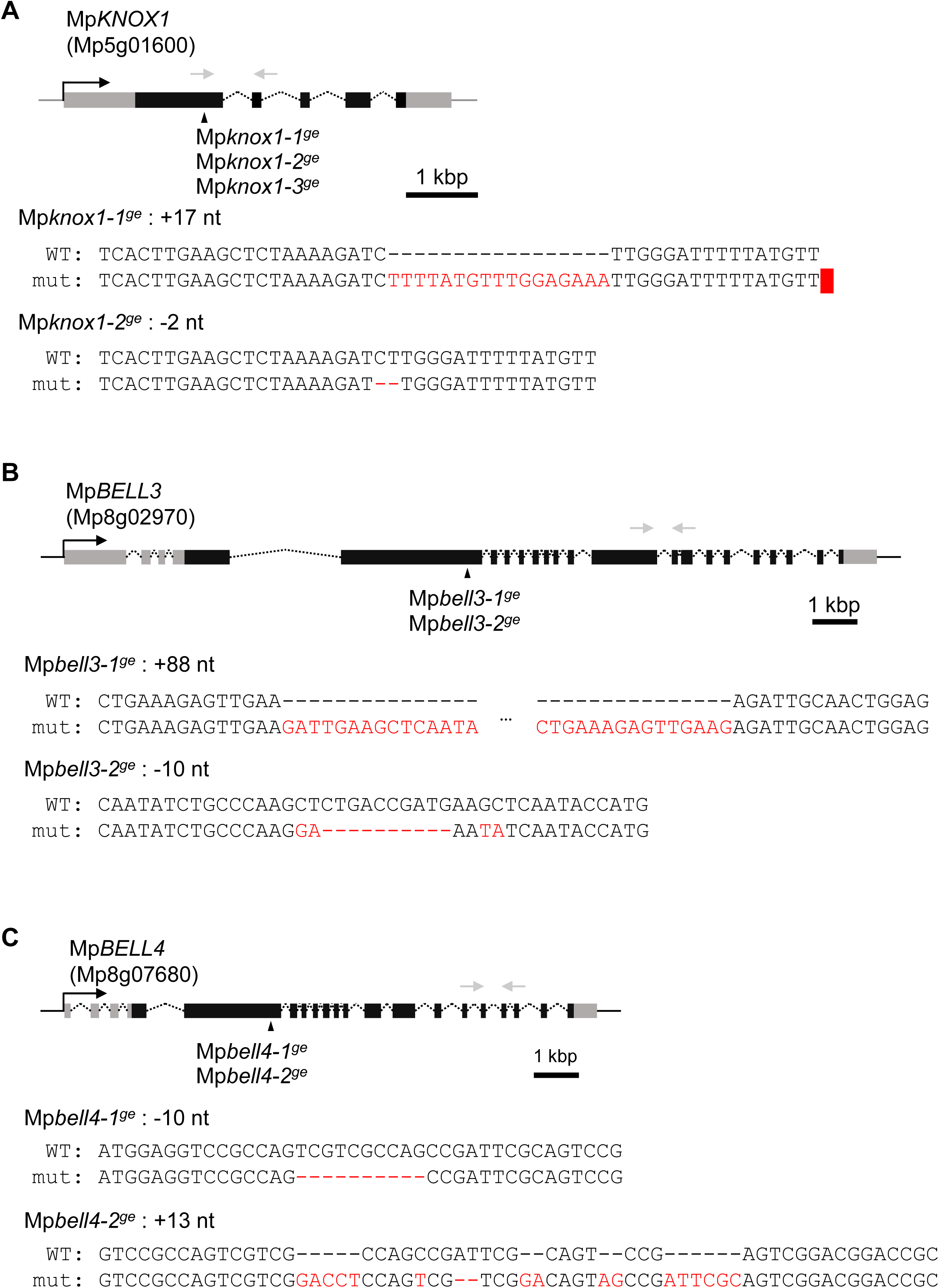
Generation of loss-of-function mutant lines by CRISPR/Cas9. Gene organization and the locations of CRISPR/Cas9-introduced mutations in Mp*KNOX1* (A), Mp*BELL3* (B) and Mp*BELL4* (C) are indicated by the following symbols; gray line, 5′- and 3′-flanking sequences; gray box, 5′- and 3′-UTR; black box, coding regions; arrowhead, mutation positions; black arrow, transcriptional direction; dotted line, splice pattern; gray arrow; primers used in RT-PCR shown in Figure 2A and 5A. Sequence alignments of wild-type and mutant alleles are shown below each gene model. Mismatched nucleotides and gaps are shown in red.

**Figure 4 - figure supplement 2.**
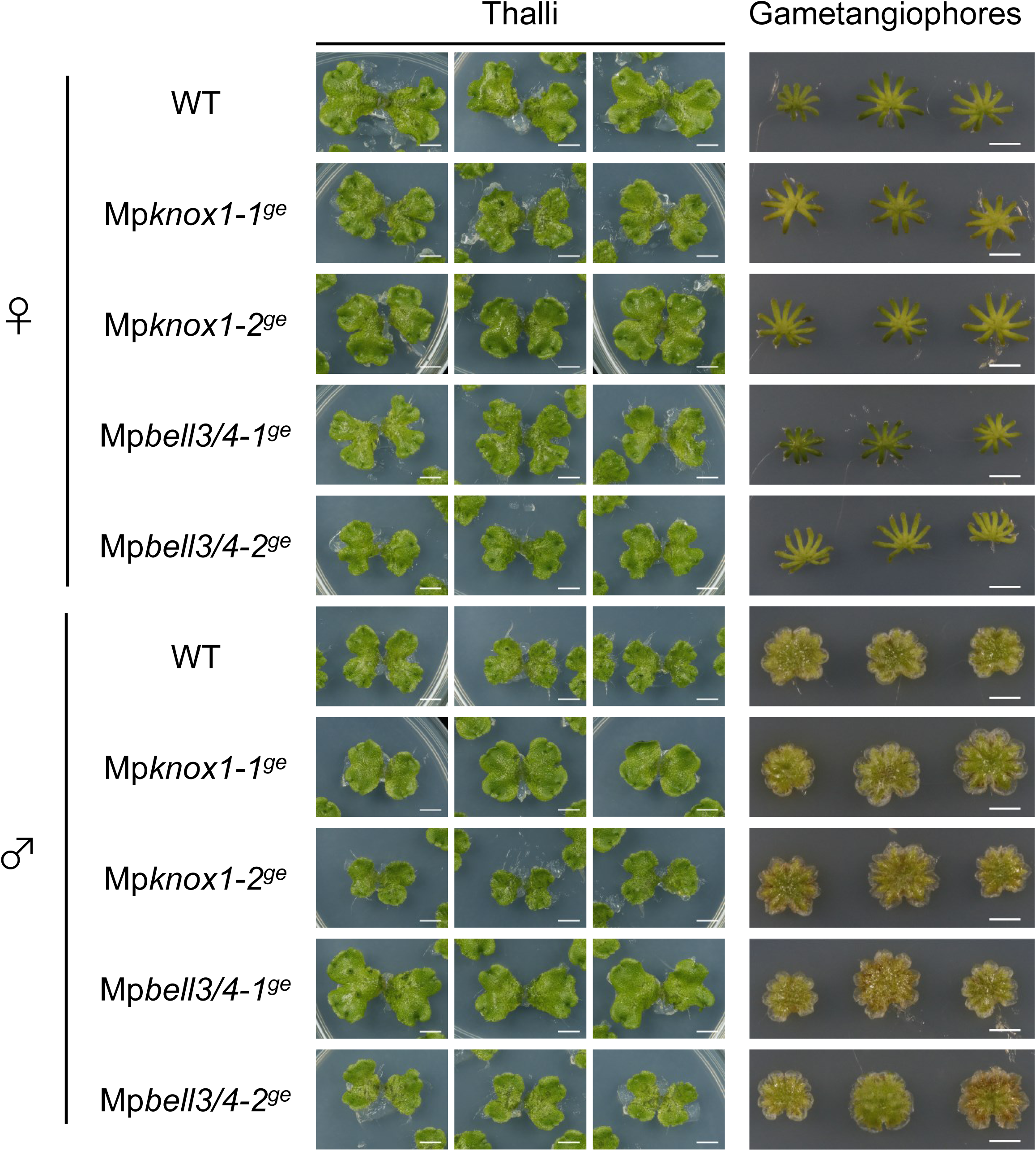
Mp*KNOX1* and Mp*BELL3/4* are dispensable for gametophyte development. Three samples of vegetative thalli and gametangiophores from wild-type (WT), Mp*knox1* and Mp*bell3/4* mutant lines are shown. Bars, 5 mm.

**Figure 4 - figure supplement 3.**
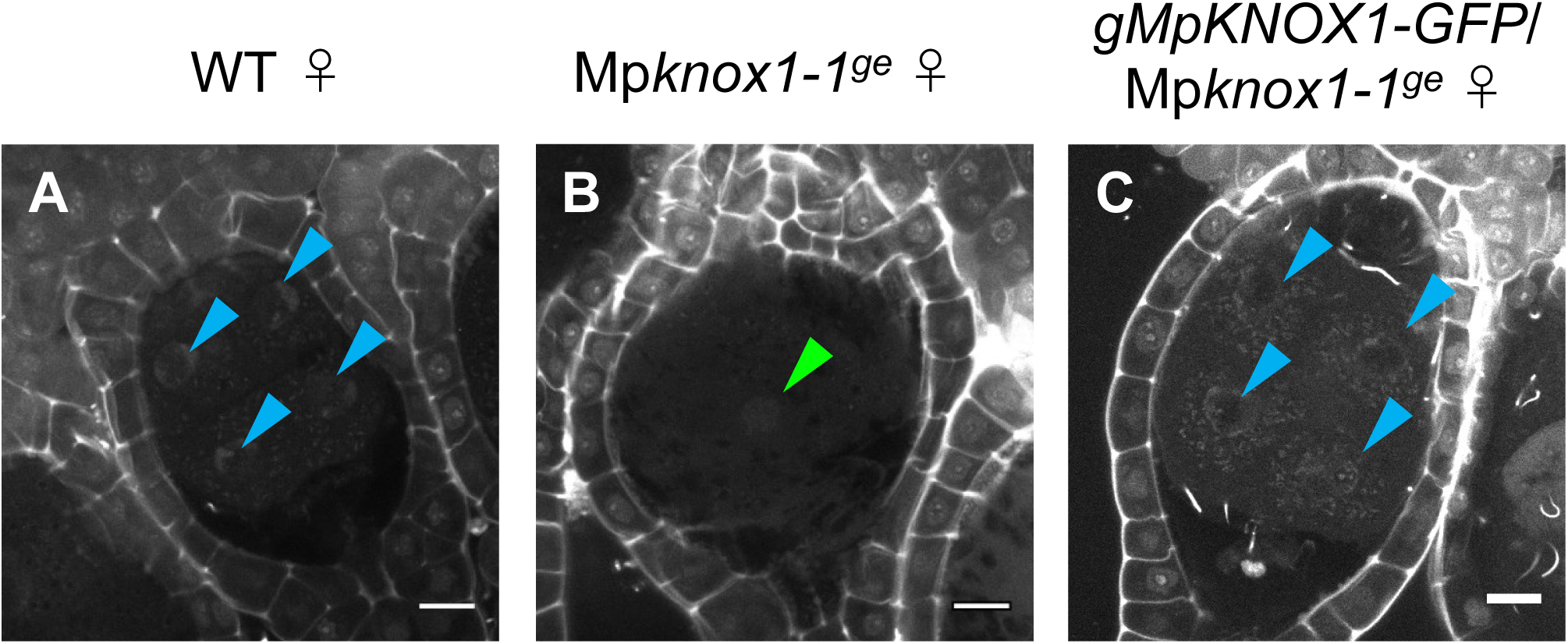
Expression of MpKNOX1-GFP driven by the Mp*KNOX1* promoter complements the karyogamy defects of Mp*knox1* mutants. 5-DAF zygotes obtained by crossing the indicated female lines with wild-type males. Green and blue arrowheads indicate a male pronucleus and embryo nuclei, respectively. Bars, 10 µm.

**Figure 4 - figure supplement 4.**
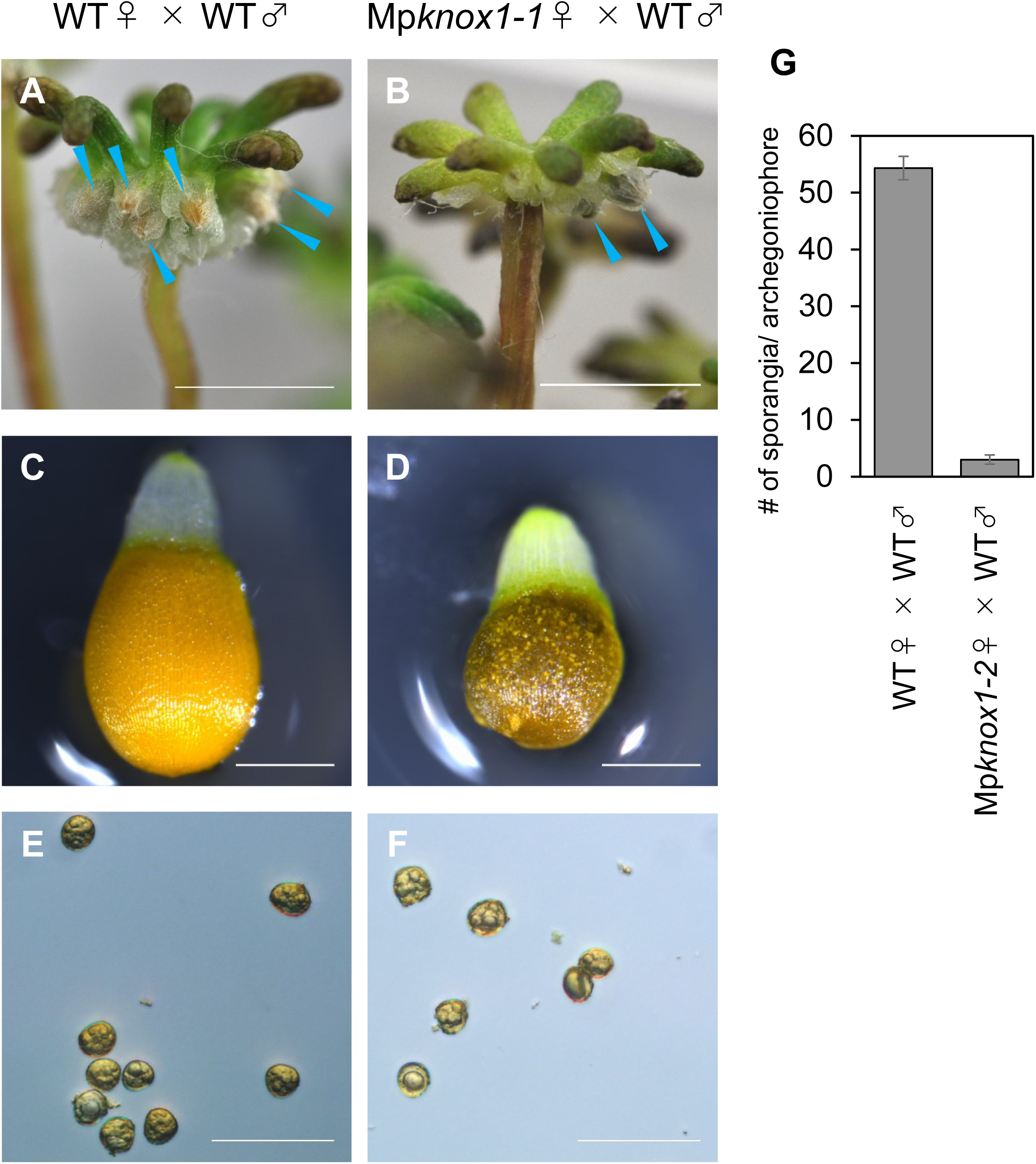
Sporangium and spore formation in wild-type and Mp*knox1* plants. **A, B.** Archegoniophores of wild-type (A) and Mp*knox1-1^ge^* (B) female plants 4 weeks after crossing with wild-type males. Blue arrowheads indicate sporangia harboring expanded capsules. **C-F.** Capsule (C, D) and spores (E, F) in wild-type (C, E) and Mp*knox1-1^ge^* (D, F) female plants crossed with wild-type males. **G.** Bar graph showing the number of sporangia per archegoniophore in wild-type and Mp*knox1-2^ge^* female plants crossed with wild-type males. Error bars indicate standard deviation. Three gametangiophores were analyzed for each genotype. Bars, 5 mm (A, B), 0.5 mm (C, D), 50 µm (E, F).

**Figure 4 - figure supplement 5.**
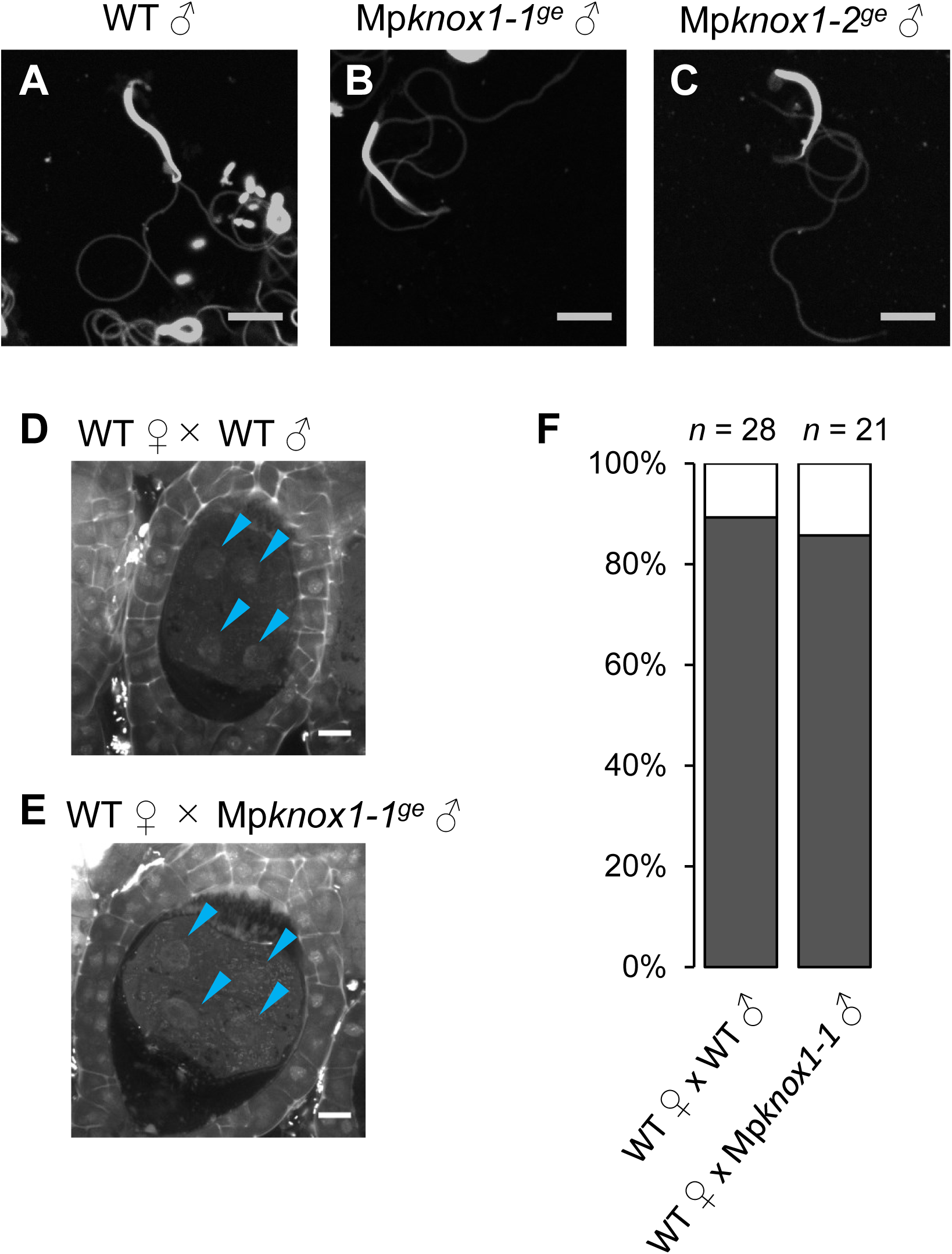
Mp*KNOX1* is dispensable for sperm differentiation and embryogenesis. **A-C.** DAPI-stained sperm cells from wild-type (A) and Mp*knox1* male plants (B, C) show indistinguishable morphology. **D, E.** 5-DAF sporophytes in wild-type female plants crossed with wild-type (C) or Mp*knox1-1^ge^* (D) males. Blue arrowheads indicate nuclei of embryo cells. **F.** Proportion of developed vs. arrested zygotes in wild-type female plants crossed with wild-type or Mp*knox1-1^ge^* males. Numbers of observed zygotes are indicated above each bar. Bars, 5 µm (A-C), 10 µm (D, E).

**Figure 5 - figure supplement 1.**
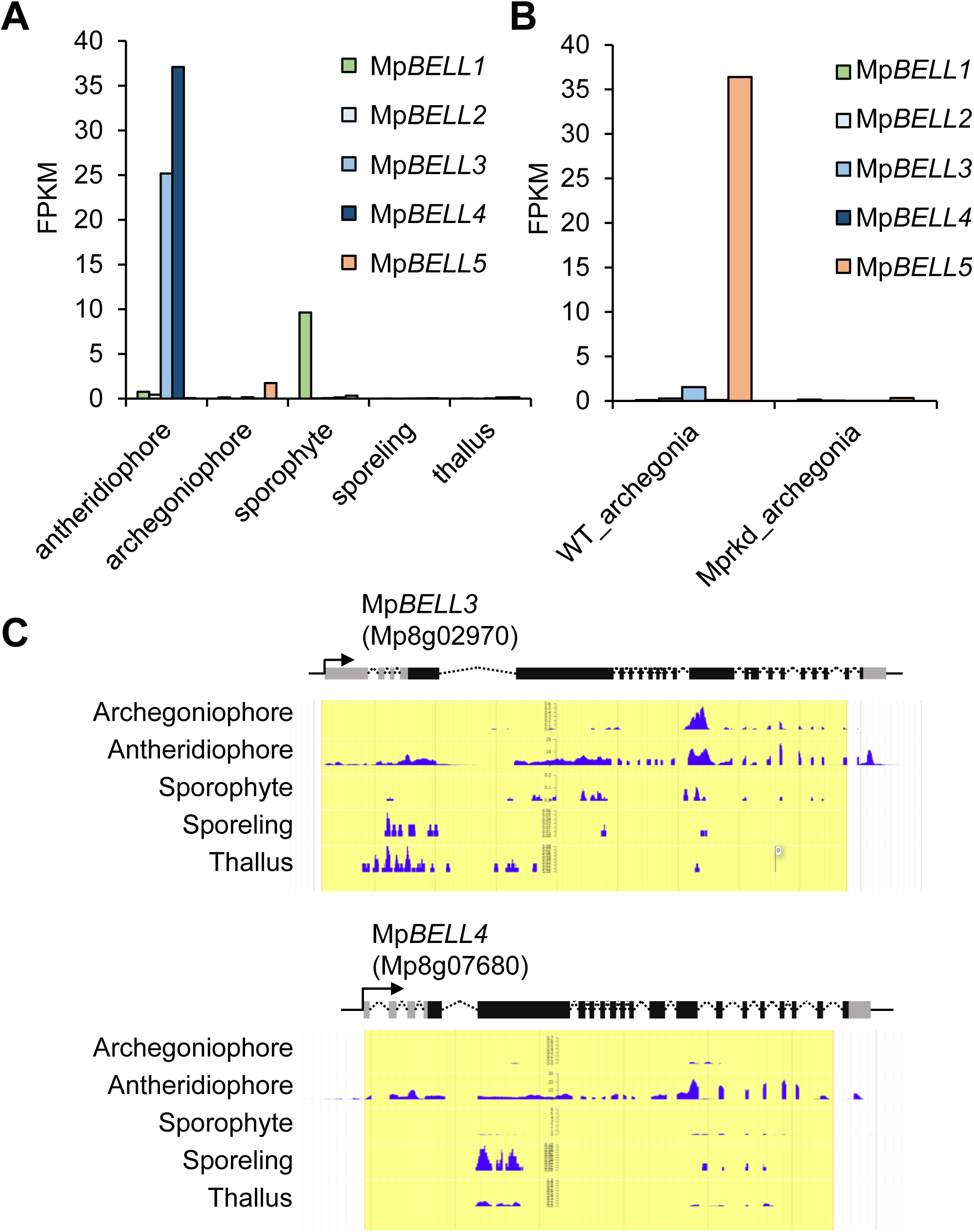
Mp*BELL3* and Mp*BELL4* are preferentially expressed in antheridiophores. **A, B.** Bar graphs showing the expression levels of Mp*BELL* genes in the indicated organs, constructed from publicly available transcriptome data (Bowman et al., 2017) (A) and the RNA-seq data obtained in this study (B). **C.** Snapshots of Genome Browser views of the Mp*BELL3* and Mp*BELL4* loci displaying their preferential expression in antheridiophores.

**Figure 5 - figure supplement 2.**
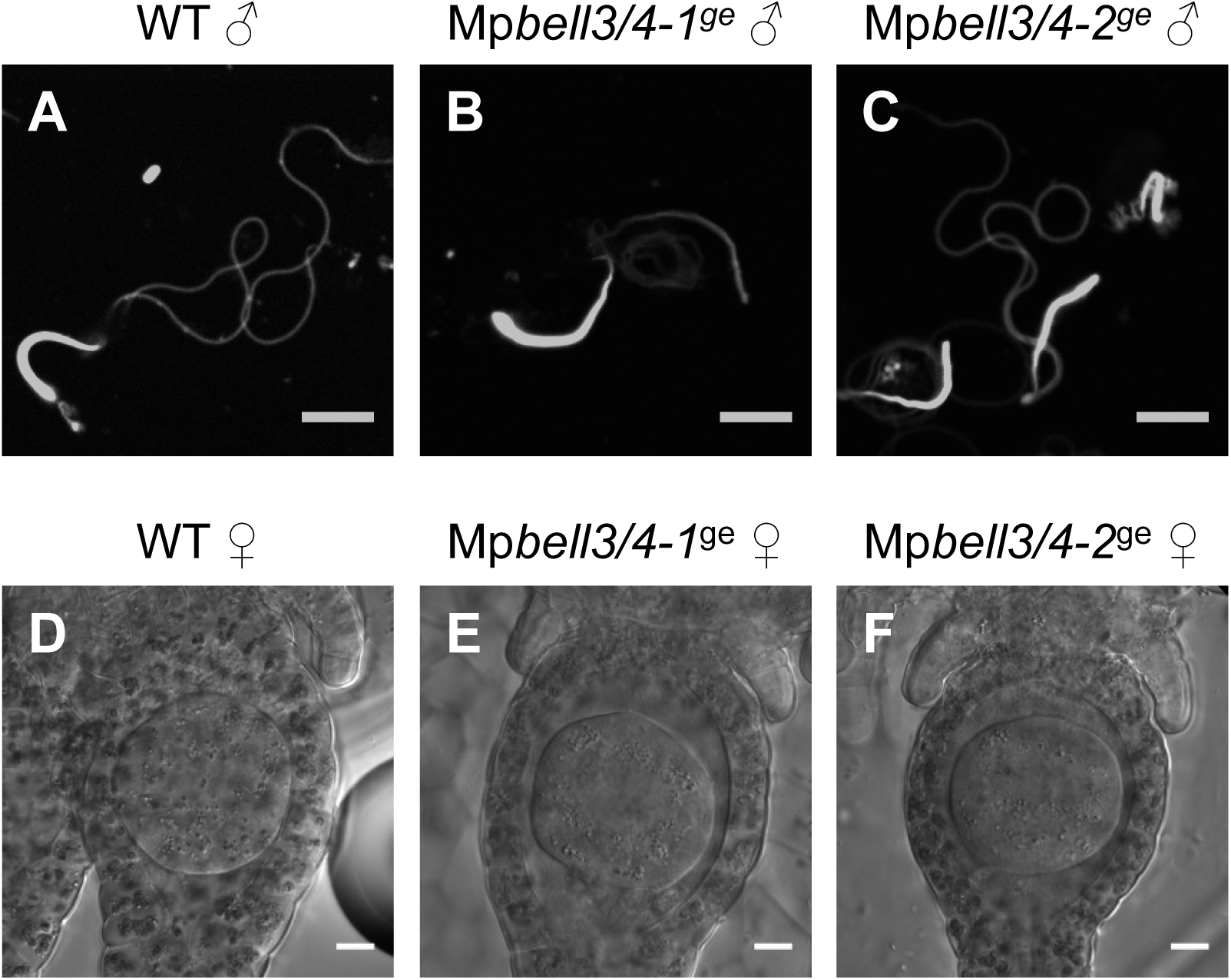
Mp*BELL3* and Mp*BELL4* are dispensable for gamete differentiation. **A-C.** DAPI-stained sperm cells from wild-type (A), Mp*bell3/4-1^ge^* (B) and Mp*bell3/4-2^ge^* (C) male plants. **D-F.** DIC images of archegonia in wild-type (D), Mp*bell3/4-1^ge^* (E) and Mp*bell3/4-2^ge^* (F) female plants. Bar, 5 µm (A-C), 10 µm (D-F).

## References

Becker, B. 2013. Snow ball earth and the split of Streptophyta and Chlorophyta. Trends in Plant Science, 18, 180–183, 10.1016/j.tplants.2012.09.010.

Bertolino, E., Reimund, B., Wildt-Perinic, D. & Clerc, R. G. 1995. A Novel Homeobox Protein Which Recognizes a Tgt Core and Functionally Interferes with a Retinoid-responsive Motif. Journal of Biological Chemistry, 270, 31178–31188, 10.1074/jbc.270.52.31178.

Bowman, J. L., Briginshaw, Liam N., And Florent, Stevie N. 2019. Evolution and co-option of developmental regulatory networks in early land plants. In: Grossniklaus, U. (ed.) Current Topics in Developmental Biology. London: Academic Press.

Bowman, J. L., Kohchi, T., Yamato, K. T., Jenkins, J., Shu, S., Ishizaki, K., Yamaoka, S., Nishihama, R., Nakamura, Y., Berger, F., Adam, C., Aki, S. S., Althoff, F., Araki, T., Arteaga-Vazquez, M. A., Balasubrmanian, S., Barry, K., Bauer, D., Boehm, C. R., Briginshaw, L., Caballero-Perez, J., Catarino, B., Chen, F., Chiyoda, S., Chovatia, M., Davies, K. M., Delmans, M., Demura, T., Dierschke, T., Dolan, L., Dorantes-Acosta, A. E., Eklund, D. M., Florent, S. N., Flores-Sandoval, E., Fujiyama, A., Fukuzawa, H., Galik, B., Grimanelli, D., Grimwood, J., Grossniklaus, U., Hamada, T., Haseloff, J., Hetherington, A. J., Higo, A., Hirakawa, Y., Hundley, H. N., Ikeda, Y., Inoue, K., Inoue, S. I., Ishida, S., Jia, Q., Kakita, M., Kanazawa, T., Kawai, Y., Kawashima, T., Kennedy, M., Kinose, K., Kinoshita, T., Kohara, Y., Koide, E., Komatsu, K., Kopischke, S., Kubo, M., Kyozuka, J., Lagercrantz, U., Lin, S. S., Lindquist, E., Lipzen, A. M., Lu, C. W., De Luna, E., Martienssen, R. A., Minamino, N., Mizutani, M., Mochizuki, N., Monte, I., Mosher, R., Nagasaki, H., Nakagami, H., Naramoto, S., Nishitani, K., Ohtani, M., Okamoto, T., Okumura, M., Phillips, J., Pollak, B., Reinders, A., R Vekamp, M., Sano, R., Sawa, S., Schmid, M. W., Shirakawa, M., Solano, R., Spunde, A., Suetsugu, N., Sugano, S., Sugiyama, A., Sun, R., Suzuki, Y., Takenaka, M., Takezawa, D., et al. 2017. Insights into land plant evolution garnered from the *Marchantia polymorpha* genome. Cell, 171, 287–304.e15, 10.1016/j.cell.2017.09.030.

Bowman, J. L., Sakakibara, K., Furumizu, C. & Dierschke, T. 2016. Evolution in the Cycles of Life. Annual Review of Genetics, 50, 133–154, 10.1146/annurev-genet-120215-035227.

De Sousa, F., Foster, P. G., Donoghue, P. C. J., Schneider, H. & Cox, C. J. 2019. Nuclear protein phylogenies support the monophyly of the three bryophyte groups (Bryophyta Schimp.). New Phytologist, 222, 565–575, 10.1111/nph.15587.

Derelle, R., Lopez, P., Guyader, H. L. & Manuel, M. 2007. Homeodomain proteins belong to the ancestral molecular toolkit of eukaryotes. Evolution & Development, 9, 212–219, 10.1111/j.1525-142X.2007.00153.x.

Durand, E. J. 1908. The development of the sexual organs and sprorogonium of Marchantia polymorpha. The Bulletin of the Torrey Botanical Club 35, 321–335, 10.2307/2485335

Fatema, U., Ali, M. F., Hu, Z., Clark, A. J. & Kawashima, T. 2019. Gamete Nuclear Migration in Animals and Plants. Frontiers in Plant Science, 10, 10.3389/fpls.2019.00517.

Ferris, P. J. & Goodenough, U. W. 1997. Mating Type in Chlamydomonas Is Specified by *mid*, the Minus-Dominance Gene. Genetics, 146, 859.

Frangedakis, E., Saint-Marcoux, D., Moody, L. A., Rabbinowitsch, E. & Langdale, J. A. 2017. Nonreciprocal complementation of Knox gene function in land plants. New Phytologist, 216, 591–604, 10.1111/nph.14318.

Furumizu, C., Alvarez, J. P., Sakakibara, K. & Bowman, J. L. 2015. Antagonistic Roles for Knox1 and Knox2 Genes in Patterning the Land Plant Body Plan Following an Ancient Gene Duplication. Plos Genetics, 11, e1004980, 10.1371/journal.pgen.1004980.

Geng, S., De Hoff, P. & Umen, J. G. 2014. Evolution of Sexes from an Ancestral Mating-Type Specification Pathway. Plos Biology, 12, e1001904, 10.1371/journal.pbio.1001904.

Geng, S., Miyagi, A. & Umen, J. G. 2018. Evolutionary divergence of the sex-determining gene *Mid* uncoupled from the transition to anisogamy in volvocine algae. Development, 145, dev162537, 10.1242/dev.162537.

Gibson, D. G., Young, L., Chuang, R. Y., Venter, J. C., Hutchison, C. A., 3rd & Smith, H. O. 2009. Enzymatic assembly of Dna molecules up to several hundred kilobases. Nat Methods, 6, 343–5, 10.1038/nmeth.1318.

Gooh, K., Ueda, M., Aruga, K., Park, J., Arata, H., Higashiyama, T. & Kurihara, D. 2015. Live-cell imaging and optical manipulation of Arabidopsis early embryogenesis. Dev Cell, 34, 242–51, 10.1016/j.devcel.2015.06.008.

Goutte, C. & Johnson, A. D. 1988. a1 protein alters the Dna binding specificity of alpha 2 repressor. Cell, 52, 875–82, 10.1016/0092-8674(88)90429-1.

Hay, A. & Tsiantis, M. 2010. Knox genes: versatile regulators of plant development and diversity. Development, 137, 3153, 10.1242/dev.030049.

Herskowitz, I. 1989. A regulatory hierarchy for cell specialization in yeast. Nature, 342, 749–57, 10.1038/342749a0.

Higo, A., Kawashima, T., Borg, M., Zhao, M., L Pez-Vidriero, I., Sakayama, H., Montgomery, S. A., Sekimoto, H., Hackenberg, D., Shimamura, M., Nishiyama, T., Sakakibara, K., Tomita, Y., Togawa, T., Kunimoto, K., Osakabe, A., Suzuki, Y., Yamato, K. T., Ishizaki, K., Nishihama, R., Kohchi, T., Franco-Zorrilla, J. M., Twell, D., Berger, F. & Araki, T. 2018. Transcription factor Duo1 generated by neo-functionalization is associated with evolution of sperm differentiation in plants. Nature Communications, 9, 5283, 10.1038/s41467-018-07728-3.

Higo, A., Niwa, M., Yamato, K. T., Yamada, L., Sawada, H., Sakamoto, T., Kurata, T., Shirakawa, M., Endo, M., Shigenobu, S., Yamaguchi, K., Ishizaki, K., Nishihama, R., Kohchi, T. & Araki, T. 2016. Transcriptional framework of male gametogenesis in the liverwort *Marchantia polymorpha* L. Plant Cell Physiol, 57, 325–38, 10.1093/pcp/pcw005.

Hisanaga, T., Okahashi, K., Yamaoka, S., Kajiwara, T., Nishihama, R., Shimamura, M., Yamato, K. T., Bowman, J. L., Kohchi, T. & Nakajima, K. 2019a. A cis-acting bidirectional transcription switch controls sexual dimorphism in the liverwort. The Embo Journal, 38, e100240, 10.15252/embj.2018100240.

Hisanaga, T., Yamaoka, S., Kawashima, T., Higo, A., Nakajima, K., Araki, T., Kohchi, T. & Berger, F. 2019b. Building new insights in plant gametogenesis from an evolutionary perspective. Nature Plants, 5, 663–669, 10.1038/s41477-019-0466-0.

Horst, N. A., Katz, A., Pereman, I., Decker, E. L., Ohad, N. & Reski, R. 2016. A single homeobox gene triggers phase transition, embryogenesis and asexual reproduction. Nature Plants, 2, 15209, 10.1038/nplants.2015.209.

Ishizaki, K., Chiyoda, S., Yamato, K. T. & Kohchi, T. 2008. *Agrobacterium*-mediated transformation of the haploid liverwort *Marchantia polymorpha* L., an emerging model for plant biology. Plant Cell Physiol, 49, 1084–91, 10.1093/pcp/pcn085.

Ishizaki, K., Nishihama, R., Ueda, M., Inoue, K., Ishida, S., Nishimura, Y., Shikanai, T. & Kohchi, T. 2015. Development of Gateway binary vector series with four different selection markers for the liverwort *Marchantia polymorpha*. PloS One, 10, e0138876, 10.1371/journal.pone.0138876.

Ishizaki, K., Nishihama, R., Yamato, K. T. & Kohchi, T. 2016. Molecular Genetic Tools and Techniques for Marchantia polymorpha Research. Plant Cell Physiol, 57, 262–70, 10.1093/pcp/pcv097.

Joo, S., Nishimura, Y., Cronmiller, E., Hong, R. H., Kariyawasam, T., Wang, M. H., Shao, N. C., El Akkad, S.-E.-D., Suzuki, T., Higashiyama, T., Jin, E. & Lee, J.-H. 2017. Gene Regulatory Networks for the Haploid-to-Diploid Transition of *Chlamydomonas reinhardtii*. Plant Physiology, 175, 314, 10.1104/pp.17.00731.

K Es, U., Richardson, W. V., Tymon, A. M., Mutasa, E. S., G Ttgens, B., Gaubatz, S., Gregoriades, A. & Casselton, L. A. 1992. The combination of dissimilar alleles of the A alpha and A beta gene complexes, whose proteins contain homeo domain motifs, determines sexual development in the mushroom Coprinus cinereus. Genes & Development, 6, 568–577.

Kariyawasam, T., Joo, S., Lee, J., Toor, D., Gao, A. F., Noh, K.-C. & Lee, J.-H. 2019. Tale homeobox heterodimer Gsm1/Gsp1 is a molecular switch that prevents unwarranted genetic recombination in Chlamydomonas. The Plant Journal, 0, 10.1111/tpj.14486.

Kerstetter, R., Vollbrecht, E., Lowe, B., Veit, B., Yamaguchi, J. & Hake, S. 1994. Sequence analysis and expression patterns divide the maize knotted1-like homeobox genes into two classes. The Plant Cell, 6, 1877, 10.1105/tpc.6.12.1877.

Koi, S., Hisanaga, T., Sato, K., Shimamura, M., Yamato, K. T., Ishizaki, K., Kohchi, T. & Nakajima, K. 2016. An evolutionarily conserved plant Rkd factor controls germ cell differentiation. Curr Biol, 26, 1775–81, 10.1016/j.cub.2016.05.013.

Kubota, A., Ishizaki, K., Hosaka, M. & Kohchi, T. 2013. Efficient *Agrobacterium*-mediated transformation of the liverwort *Marchantia polymorpha* using regenerating thalli. Biosci Biotechnol Biochem, 77, 167–72, 10.1271/bbb.120700.

Kurihara, D., Mizuta, Y., Sato, Y. & Higashiyama, T. 2015. ClearSee: a rapid optical clearing reagent for whole-plant fluorescence imaging. Development, 142, 4168–79, 10.1242/dev.127613.

Lee, J.-H., Lin, H., Joo, S. & Goodenough, U. 2008. Early Sexual Origins of Homeoprotein Heterodimerization and Evolution of the Plant Knox/Bell Family. Cell, 133, 829–840, https://doi.org/10.1016/j.cell.2008.04.028.

Ligrone, R., Duckett, J. G. & Renzaglia, K. S. 2012a. Major transitions in the evolution of early land plants: a bryological perspective. Annals of Botany, 109, 851–871, 10.1093/aob/mcs017.

Ligrone, R., Duckett, J. G. & Renzaglia, K. S. 2012b. The origin of the sporophyte shoot in land plants: a bryological perspective. Annals of Botany, 110, 935–941, 10.1093/aob/mcs176.

Lopez, D., Hamaji, T., Kropat, J., De Hoff, P., Morselli, M., Rubbi, L., Fitz-Gibbon, S., Gallaher, S. D., Merchant, S. S., Umen, J. & Pellegrini, M. 2015. Dynamic Changes in the Transcriptome and Methylome of *Chlamydomonas reinhardtii* throughout Its Life Cycle. Plant Physiology, 169, 2730, 10.1104/pp.15.00861.

Love, M. I., Huber, W. & Anders, S. 2014. Moderated estimation of fold change and dispersion for Rna-seq data with Deseq2. Genome Biology, 15, 550, 10.1186/s13059-014-0550-8.

Luo, A., Shi, C., Zhang, L. & Sun, M.-X. 2014. The expression and roles of parent-of-origin genes in early embryogenesis of angiosperms. Frontiers in Plant Science, 5, 10.3389/fpls.2014.00729.

Miyashima, S., Roszak, P., Sevilem, I., Toyokura, K., Blob, B., Heo, J. O., Mellor, N., Help-Rinta-Rahko, H., Otero, S., Smet, W., Boekschoten, M., Hooiveld, G., Hashimoto, K., Smetana, O., Siligato, R., Wallner, E. S., Mahonen, A. P., Kondo, Y., Melnyk, C. W., Greb, T., Nakajima, K., Sozzani, R., Bishopp, A., De Rybel, B. & Helariutta, Y. 2019. Mobile Pear transcription factors integrate positional cues to prime cambial growth. Nature, 565, 490–494, 10.1038/s41586-018-0839-y.

Motohashi, K. 2015. A simple and efficient seamless Dna cloning method using Slice from *Escherichia coli* laboratory strains and its application to SliP site-directed mutagenesis. Bmc Biotechnol, 15, 47, 10.1186/s12896-015-0162-8.

Mukherjee, K., Brocchieri, L. & B Rglin, T. R. 2009. A Comprehensive Classification and Evolutionary Analysis of Plant Homeobox Genes. Molecular Biology and Evolution, 26, 2775–2794, 10.1093/molbev/msp201.

Ning, J., Otto, T. D., Pfander, C., Schwach, F., Brochet, M., Bushell, E., Goulding, D., Sanders, M., Lefebvre, P. A., Pei, J., Grishin, N. V., Vanderlaan, G., Billker, O. & Snell, W. J. 2013. Comparative genomics in Chlamydomonas and Plasmodium identifies an ancient nuclear envelope protein family essential for sexual reproduction in protists, fungi, plants, and vertebrates. Genes & Development, 27, 1198–1215.

Nishimura, Y., Shikanai, T., Nakamura, S., Kawai-Yamada, M. & Uchimiya, H. 2012. Gsp1 Triggers the Sexual Developmental Program Including Inheritance of Chloroplast Dna and Mitochondrial Dna in *Chlamydomonas reinhardtii*. The Plant Cell, 24, 2401, 10.1105/tpc.112.097865.

Nozaki, H., Mori, T., Misumi, O., Matsunaga, S. & Kuroiwa, T. 2006. Males evolved from the dominant isogametic mating type. Current Biology, 16, R1018–R1020, 10.1016/j.cub.2006.11.019.

Parker, G. A., Baker, R. R. & Smith, V. G. F. 1972. The origin and evolution of gamete dimorphism and the male-female phenomenon. Journal of Theoretical Biology, 36, 529–553, https://doi.org/10.1016/0022-5193(72)90007-0.

Puttick, M. N., Morris, J. L., Williams, T. A., Cox, C. J., Edwards, D., Kenrick, P., Pressel, S., Wellman, C. H., Schneider, H., Pisani, D. & Donoghue, P. C. J. 2018. The Interrelationships of Land Plants and the Nature of the Ancestral Embryophyte. Current Biology, 28, 733–745.e2, 10.1016/j.cub.2018.01.063.

R Vekamp, M., Bowman, J. L. & Grossniklaus, U. 2016. *Marchantia* Mprkd regulates the gametophyte-sporophyte transition by keeping egg cells quiescent in the absence of fertilization. Curr Biol, 26, 1782–9, 10.1016/j.cub.2016.05.028.

Sakakibara, K., Ando, S., Yip, H. K., Tamada, Y., Hiwatashi, Y., Murata, T., Deguchi, H., Hasebe, M. & Bowman, J. L. 2013. Knox2 Genes Regulate the Haploid-to-Diploid Morphological Transition in Land Plants. Science, 339, 1067, 10.1126/science.1230082.

Sakakibara, K., Nishiyama, T., Deguchi, H. & Hasebe, M. 2008. Class 1 Knox genes are not involved in shoot development in the moss Physcomitrella patens but do function in sporophyte development. Evolution & Development, 10, 555–566, 10.1111/j.1525-142X.2008.00271.x.

Shimamura, M. 2016. *Marchantia polymorpha*: taxonomy, phylogeny and morphology of a model system. Plant Cell Physiol, 57, 230–56, 10.1093/pcp/pcv192.

Spit, A., Hyland, R. H., Mellor, E. J. C. & Casselton, L. A. 1998. A role for heterodimerization in nuclear localization of a homeodomain protein. Proceedings of the National Academy of Sciences, 95, 6228, 10.1073/pnas.95.11.6228.

Sugano, S. S., Nishihama, R., Shirakawa, M., Takagi, J., Matsuda, Y., Ishida, S., Shimada, T., Hara-Nishimura, I., Osakabe, K. & Kohchi, T. 2018. Efficient Crispr/Cas9-based genome editing and its application to conditional genetic analysis in *Marchantia polymorpha*. PloS One, 13, e0205117, 10.1101/277350.

Sun, J., Nishiyama, T., Shimizu, K. & Kadota, K. 2013. Tcc: an R package for comparing tag count data with robust normalization strategies. Bmc Bioinformatics, 14, 219, 10.1186/1471-2105-14-219.

Trapnell, C., Pachter, L. & Salzberg, S. L. 2009. TopHat: discovering splice junctions with Rna-Seq. Bioinformatics, 25, 1105–11, 10.1093/bioinformatics/btp120.

Tsuboyama, S., Nonaka, S., Ezura, H. & Kodama, Y. 2018. Improved G-AgarTrap: A highly efficient transformation method for intact gemmalings of the liverwort Marchantia polymorpha. Scientific Reports, 8, 10800, 10.1038/s41598-018-28947-0.

Uchiumi, T., Uemura, I. & Okamoto, T. 2007. Establishment of an in vitro fertilization system in rice (Oryza sativa L.). Planta, 226, 581–9, 10.1007/s00425-007-0506-2.

Yamaoka, S., Nishihama, R., Yoshitake, Y., Ishida, S., Inoue, K., Saito, M., Okahashi, K., Bao, H., Nishida, H., Yamaguchi, K., Shigenobu, S., Ishizaki, K., Yamato, K. T. & Kohchi, T. 2018. Generative cell specification requires transcription factors evolutionarily conserved in land plants. Curr Biol, 28, 479–486.e5, 10.1016/j.cub.2017.12.053.

Zinsmeister, D. D. & Carothers, Z. B. 1974. The fine structure of oogenesis in *Marchantia polymorpha*. American Journal of Botany, 61, 499–512, 10.1002/j.1537-2197.1974.tb10789.x.

